# Intracellular *Pseudomonas aeruginosa* persist and evade antibiotic treatment in a wound infection model

**DOI:** 10.1101/2024.05.15.594279

**Authors:** Stéphane Pont, Flore Nilly, Laurence Berry, Anne Bonhoure, Morgan A. Alford, Mélissande Louis, Pauline Nogaret, Olivier Lesouhaitier, Robert E. W. Hancock, Patrick Plésiat, Anne-Béatrice Blanc-Potard

## Abstract

Persistent bacterial infections evade host immunity and resist antibiotic treatments through various mechanisms that are difficult to evaluate in a living host. *Pseudomonas aeruginosa* is a main cause of chronic infections in patients with cystic fibrosis (CF) and wounds. Here, by immersing wounded zebrafish embryos in a suspension of *P. aeruginosa* isolates from CF patients, we established a model of persistent infection that mimics a murine chronic skin infection model. Live and electron microscopy revealed persisting aggregated *P. aeruginosa* inside zebrafish cells, including macrophages, at unprecedented resolution. Persistent *P. aeruginosa* exhibited adaptive resistance to several antibiotics, host cell permeable drugs being the most efficient. Moreover, persistent bacteria could be partly re-sensitized to antibiotics upon addition of anti-biofilm molecules that dispersed the bacterial aggregates *in vivo*. Collectively, this study demonstrates that an intracellular location protects *P. aeruginosa in vivo* from host innate immunity and antibiotics, and provides new insights into efficient treatments against chronic infections.

## Introduction

Many bacterial pathogens can persist inside their hosts for long periods of time, due to immunodeficiency of the host, immune evasion by the bacteria and/or ineffective antibiotic treatments. *Pseudomonas aeruginosa* is a Gram-negative bacterium ubiquitous in watery environments and a major cause of a variety of nosocomial infections worldwide ^1^, including urinary tract, respiratory tract, wound/skin and blood infections. *P. aeruginosa* infections can be either acute or chronic, implying different virulence strategies to evade host immunity ^1^. Chronic colonization is frequent during cystic fibrosis (CF), non-CF bronchiectasis or chronic obstructive pulmonary disease (COPD), and also occurs on wounds ^2,3^. In long-lasting lung infections in CF patients, *P. aeruginosa* adopts a sessile lifestyle, with bacterial communities organized in a biofilm structure, impeding bacterial clearance by the immune system ^4^.

*P. aeruginosa*, which is well known for its capacity to develop resistance to antibiotic treatments, belongs to the threatening group of ESKAPEE pathogens, and is recognized by the World Health Organization as a critical priority for new therapeutics ^5^. Chronic bacterial infections are challenging to treat with antibiotics due to a combination of intrinsic, acquired, and adaptive drug resistance, along with the specific bacterial lifestyle in infected tissues. The adaptive resistance to antibiotics, which is a phenotypic non-heritable trait ^6^, is related to several factors, including biofilm lifestyle, slow growth and low metabolic activity ^7^. Moreover, increasing evidence supports the idea that *P. aeruginosa* undergoes an intracellular life cycle during infection ^8,9^, including human pulmonary infection as shown with the recent observation of intracellular *P. aeruginosa* in the airway epithelium of CF lung explants ^10^. Whereas intracellular localization was shown to affect drug efficacy against *P. aeruginosa* in cell culture models ^11-13^, this aspect has not been addressed *in vivo*, especially in the context of a persistent infection.

While a plethora of *in vivo* models have been used to assess *P. aeruginosa* virulence ^14^, only a small subset were aimed at studying this pathogen in the context of persistent colonization and testing the efficacy of treatments on chronic infection. Current animal models to study *P. aeruginosa* respiratory chronic pathogenesis mainly rely on lung administration of bacteria embedded in agar/agarose beads ^15^. A murine chronic skin infection model has also been developed, mimicking long-term colonization in humans ^16^. Zebrafish (*Danio rerio*), a vertebrate that shows the advantages of invertebrates (moderate ethical issues, low cost, high production of eggs), represents an appealing *in vivo* model for drug testing and high-resolution real-time visualization of bacteria and host cells due to embryo transparency. More specifically, the zebrafish innate immune system is very similar to that of mammals, notably regarding phagocytic cells (neutrophils and macrophages) and soluble immune mediators such as cytokines and complement proteins ^17^. Real time *in vivo* imaging has previously allowed the visualization of the interaction of *P. aeruginosa* with innate immune cells in the context of acute infections ^18-20^.

In this study, we established for the first time a model of persistent colonization of the zebrafish embryo by *P. aeruginosa*, using a caudal wound infection protocol and *P. aeruginosa* CF clinical isolates. The visualization of persisting bacteria revealed the importance of an intracellular phase, notably inside macrophages, and the formation of bacterial aggregates *in vivo*. Persistent bacteria became less responsive to treatment by several classes of antibiotics, and host cell permeable antibiotics were found to be more efficient. In addition, antibiotic efficacy against persistent bacteria could be potentiated by addition of anti-biofilm compounds. This work offers an *in vivo* model to investigate the contribution of intracellular *P. aeruginosa* to chronic infection, with unprecedented imaging capabilities, and paves the way to assess the efficacy of therapeutics in the context of a persistent colonization.

## Results

### Zebrafish embryo is an appropriate model to monitor a persistent infection with *P. aeruginosa* CF isolates

While reference laboratory strains causing acute infections (PAO1, PA14, and PAK) have been widely used in the zebrafish model to gain insights into the host-*P. aeruginosa* interaction *in vivo*, investigations of the pathogenesis of clinical isolates in this vertebrate model have remained very scarce, and relied on the microinjection of single CF isolates ^21,22^. We recently developed a wound infection protocol in zebrafish based on the immersion of tail fin-amputated embryos with *P. aeruginosa* PAO1 strain, which caused an acute infection within 20 hpi ^23^. Here, we used this infection mode to evaluate *in vivo* the virulence of three CF isolates (A6520, B6513 and C6490), that were not tested earlier in any infection model, to ensure an unbiased approach. The survival of zebrafish embryos was first monitored over 40 hours post infection (hpi) (Figure 1A). All three CF isolates were highly attenuated, being associated with 90% (A6520, B6513) to 100% (C6490) survival, when compared to the reference strain PAO1 (50% survival) (Figure 1B). Additional experiments were next conducted on these CF isolates to assess whether their attenuated phenotype was linked to the elimination or not of the bacteria by the host.

**Figure 1.**
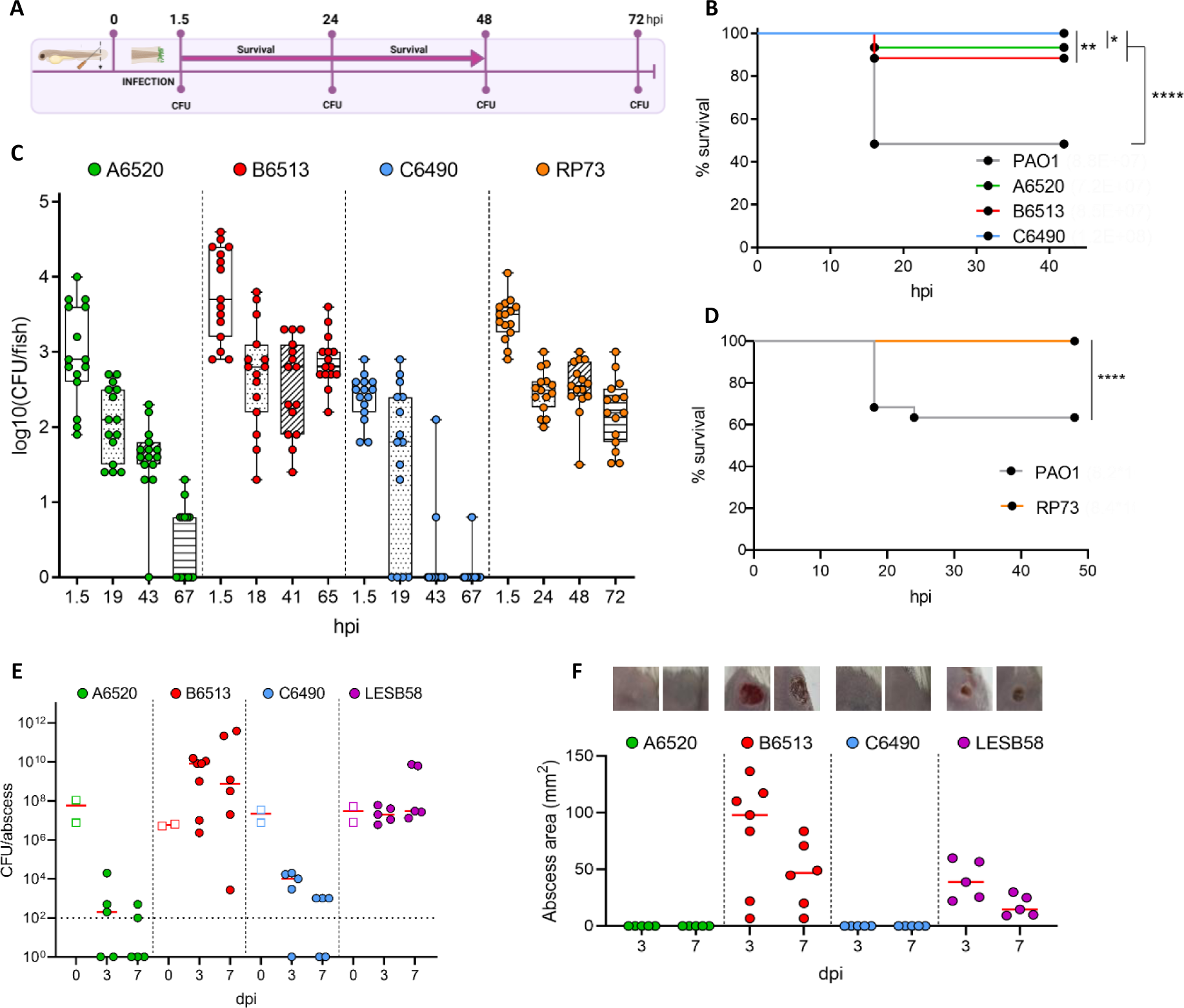
CF isolates can establish a persistent infection in zebrafish embryos, with parallel profiles to a murine model of infection. (A) Experimental timeline to assess bacterial virulence and persistence in zebrafish embryo. (B) and (D) Embryo survival following infection by immersion with indicated *P. aeruginosa* strains at bacterial concentrations ranging from 7.2x10^7^ to 1.2x10^8^ CFU/mL. Survival was monitored for >40h following the infection (n=3, 60 larvae). Log-rank test: * *P*<0.05, ** *P*<0.01 and **** *P*<0.0001. (C) Evolution of the bacterial load per embryo over time (until 72 hpi). Following infection by immersion with GFP^+^ *P. aeruginosa* clinical isolates, embryos were crushed at the indicated time points and plated for CFU counting (n=3, 15 larvae). (E) and (F) Virulence and persistence of CF isolates in a murine model of high-density cutaneous infection. Bacterial load (E) and size of abscesses (F) formed for three or seven days in CD-1 mice subcutaneously injected with indicated *P. aeruginosa* strains in the right dorsum were quantified. The CFUs reported at 0 dpi (empty squares) correspond to the injected inoculum. At 3 dpi and 7 dpi, abscesses were measured and harvested in phosphate buffered saline (PBS), homogenized and plated on lysogeny broth (LB) for bacterial enumeration. Data from two independent experiments containing 2–4 biological replicates each (*n* = 5-7) are displayed as the median. The limit of detection (LOD) is displayed as a dashed line at 10^2^ CFU/abscess. The photo insets above the graph are representative images from treatment groups.

To discriminate between bacterial clearance and bacterial persistence, we assessed the evolution of the bacterial load per embryo over 72 hpi. To easily recognize and count *P. aeruginosa* bacteria, we used green-fluorescent strains harboring a chromosomally-integrated *gfp* gene with constitutive expression, to avoid the risk of GFP signal loss upon time. For all strains, a notable variability between individual larvae was noticed, with some differences being >100-fold (2 log_10_). With isolates A6520 and C6490, the majority of the embryos were observed to be free of bacteria at 67 or 43 hpi, respectively (Figure 1C). Conversely, with strain B6513, 90% of the bacteria were eliminated between 1.5 to 18 hpi, but the remaining bacterial load (median of 630 CFU) stayed relatively constant in numbers until 65 hpi (Figure 1C). Thus, isolate B6513 developed a persistent infection in this immersion zebrafish model, whereby a subset of bacteria survived after 18 hpi without killing the host. The residual CFU analysis for isolate B6513 was extended once up to 6 days post-infection (dpi) and a stable number of persistent bacteria per fish was observed. Subsequent experiments were carried out only on embryos for up to 3 dpi (i.e. 5 days post-fertilization, within the frame not regulated as animal experiments) to follow the 3R-principle and limit a possible reinfection upon food ingestion.

To extend the relevance of this model, we assessed the virulence and persistence of the well characterized isolate RP73, a late CF isolate capable of long-term colonization in mouse airways ^24,25^. Consistent with previous mouse studies, and similarly to strain B6513, the RP73 isolate failed to induce embryo mortality and persisted in infected zebrafish embryos (Figures 1C and 1D). To verify that the persistent phenotype reflected colonization of the wound and not the ability of the isolates to adhere to the embryo’s skin, we assessed the bacterial load over time in uninjured embryos incubated with persistent strains B6513 and RP73. At 24 hpi, no CFU were detected for either isolate (Figure S1), showing that the injury was required to drive a persistent infection.

We next addressed the behaviour of the three CF isolates A6520, B6513 and C6490 in a mouse model of chronic infection that involves cutaneous abscesses ^16^. *P. aeruginosa* LESB58, a well-characterized CF isolate that causes chronic lung infection and chronic skin abscess in mice ^16,26^, was used as a reference to assess *in vivo* growth and virulence of the clinical isolates over a seven-day period after subcutaneous injection in mice (with approx. 5 x 10^7^ CFU). As previously reported, *P. aeruginosa* LESB58 persisted at the infection site, being recovered at median densities above 10^7^ CFU/abscess at 3 and 7 dpi (Figure 1E). Similarly, the clinical isolate *P. aeruginosa* B6513 was recovered at high median densities, >10^8^ CFU/abscess at 3 and 7 dpi. In contrast, clinical isolates *P. aeruginosa* A6520 and C6490 were mostly eliminated from the host when compared to the other strains, as reflected by low median bacterial densities (< 10^4^ CFU/abscess at 3 dpi, and 10^3^ or below the limit of detection at 7 dpi). Trends in abscess sizes were reflective of bacterial recovery from the localized infection site at both time points, with abscess (dermonecrotic cutaneous tissue) lesion areas of 15-39 mm^2^ and 47-98 mm^2^ for strains LESB58 and B6513, respectively, whereas strains A6520 and C6490 did not cause abscess formation (Figure 1F). Healing of the cutaneous tissue appeared to occur at 7 dpi for both *P. aeruginosa* LESB58 and B6513, since the sizes of abscesses had decreased (-24 and -51 mm^2^ respectively) relative to the abscesses of animals sacrificed 3 dpi. Accordingly, the mean clinical scores of animals decreased during the time from 3 to 7 dpi across treatment groups (Figure S2). Furthermore, clinical scores of animals remained low overall, with a maximum recorded total score of 8/50 (in the *P. aeruginosa* B6513 treatment group), indicating that none of the strains caused severe disease in animals.

Taken together, our results showed that the injured embryo immersion model of zebrafish infection is suitable to assess persistence of *P. aeruginosa* clinical isolates. Importantly, the persistence profiles of strains in zebrafish are consistent with the phenotypes observed in a murine chronic infection model. We next took advantage of the unique imaging opportunities offered by the zebrafish embryo to visualize persisting bacteria at high resolution.

### Dynamic interaction of persistent *P. aeruginosa* with host immune cells *in vivo*

Macrophages and neutrophils have been shown to rapidly phagocytose *P. aeruginosa* upon zebrafish infection ^20,27-29^. We took advantage of the optical transparency of embryos to image GFP-expressing isolate B6513 throughout the infection, assess bacterial morphology, localization and interaction with recruited macrophages, using Tg*(mfap4::mCherry-F)* zebrafish embryos that harbor red fluorescent macrophages. Consistent with CFU counts, we observed an important decrease in the bacterial load between 1.5 and 24 hpi (Figure 2A). At 1.5 hpi, most bacteria appeared free living, while at 24 and 72 hpi, they mainly appeared as aggregates localized nearby the injury site (Figure 2A). Intra-macrophagic *P. aeruginosa* were present throughout the course of infection (Figures 2A and 2B). The presence of bacterial aggregates, with putative localization inside macrophages, was not specific to the B6513 isolate since similar findings were visualized with the RP73 strain (Figure S3). Bacterial aggregates were retained within macrophages for up to 10 h, as shown using time-lapse confocal microscopy (Figure 2C). The size distribution of GFP positive spots was stable over time, with a volume mainly comprised between 1.5 µm^3^ (individual bacteria) and 100 µm^3^ (clusters) in size (Figure 2D). Larger aggregates (>100 µm^3^) were observed in some embryos. Infection seemed to delay macrophage recruitment to the wound at the early time point (1.5 hpi, Figure 2E). Macrophage recruitment in infected embryos peaked at 24 hpi and decreased slowly, being close to the level of uninfected controls at 72 hpi (Figures 2E and S4).

**Figure 2.**
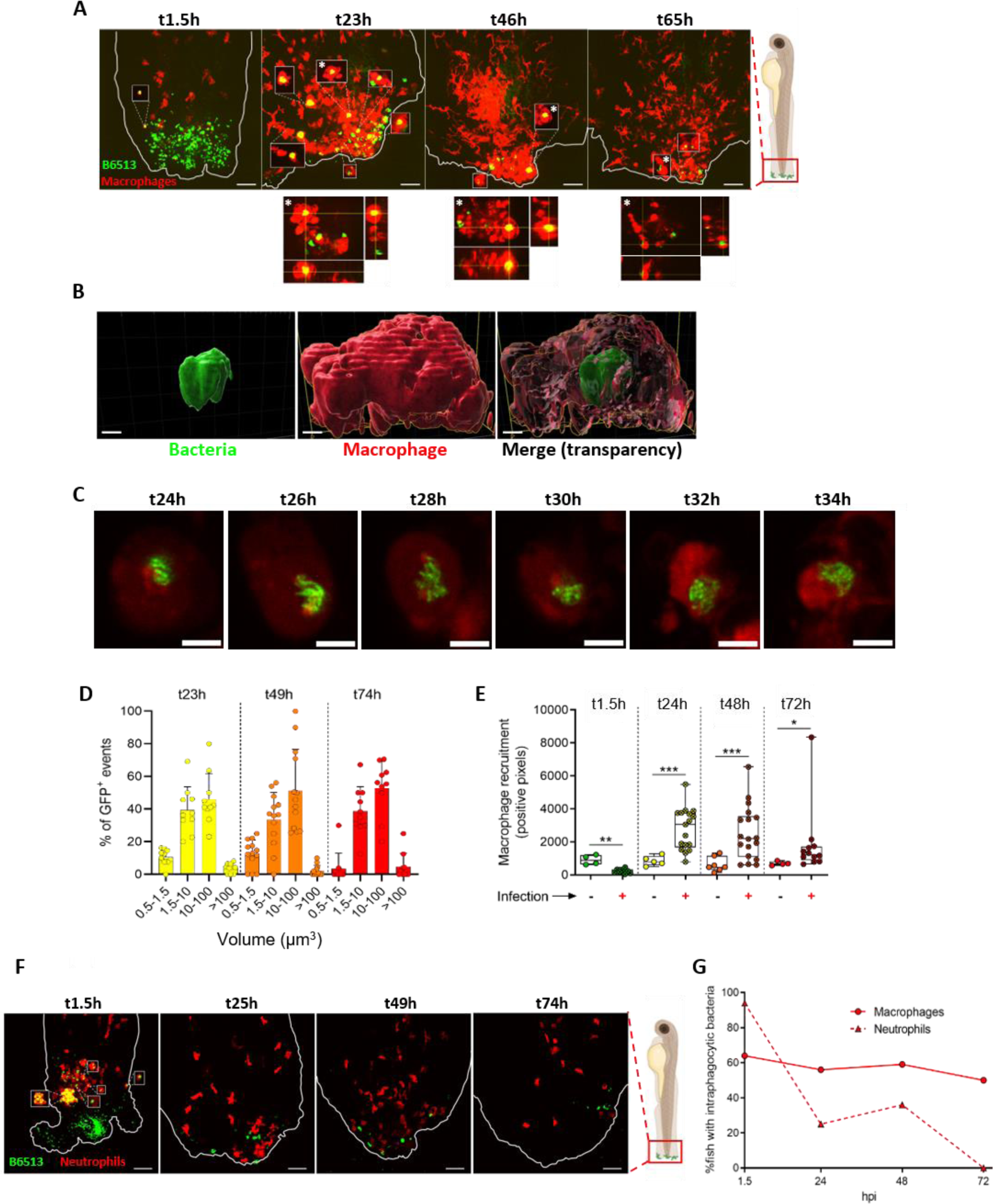
Upon persistent colonization, *P. aeruginosa* B6513 form aggregates which can be visualized inside macrophages but not neutrophils. (A) Representative maximal projections of confocal images, showing interactions between bacteria (green) and recruited macrophages (red) in Tg(*mfap4:mCherry-F*) larvae at different time points. White rectangles denote images extracted from a single optical section of macrophages with intracellular *P. aeruginosa*. Below the images, orthogonal representations of the (*) events are shown, confirming that bacteria were inside macrophages. Scale bar: 40 µm. Note that pictures come from different embryos imaged for each indicated times. (B) 3D reconstruction allowing to confirm the intramacrophagic localization of a bacterial aggregate. Scale bar: 7 µm. (C) Zoomed single optical sections of an intramacrophagic bacterial aggregate tracked from 24 hpi to 34 hpi using time-lapse confocal imaging. Image acquisition was done every 30 min and images corresponding to 2 h intervals are shown. Brightness/contrast settings were modified comparatively to (A) for a better visualization of the aggregate organization after a deconvolution. Scale bar: 5 µm. (D) Volume repartition of GFP^+^ events per larvae quantified following 3D reconstruction (10 to 13 embryos were imaged at each time point). (E) Macrophage quantification at the wound, measured by the number of red pixels at the median plan of the stack, in presence or absence of bacteria at various time points (4 to 8 control and 8 to 18 infected embryos were imaged at each time point). Mann-Whitney test: \**P*<0.05, \*\**P*<0.01 and \*\*\**P*<0.001. (F) Representative maximal projections of confocal images, showing interactions between bacteria (green) and recruited neutrophils (red) in Tg(*LysC*:*dsRed*) larvae. White rectangles denote images extracted from a single optical section of neutrophils with intracellular *P. aeruginosa*. Scale bar: 40 µm. Note that pictures come from different embryos imaged for each indicated times. (G) Evolution of the proportion of Tg(*mfap4:mCherry-F*) and Tg(*LysC*:*dsRed*) larvae with at least one event of intra-macrophage or intra-neutrophil bacteria, respectively, over time (10 to 16 larvae were imaged at each time point).

The interaction of persistent bacteria with neutrophils using the Tg(*LysC*:*dsRed*) transgenic line harboring red fluorescent neutrophils was subsequently investigated (Figure S5). In agreement with previous experiments done on microinjected embryos ^27-29^, numerous intra-neutrophilic *P. aeruginosa* were observed shortly after colonization (Figures 2F and 2G). However, this frequency was strongly reduced at 24 and 48 hpi, and no intra-neutrophil bacteria were observed at 72 hpi. Thus, the kinetics of appearance of intra-neutrophil and intra-macrophage bacteria were very different (Figure 2G), suggesting that the interaction with neutrophils was restricted to the early steps of colonization and/or that the killing activity of neutrophils and/or their short half-life prevented any intra-neutrophilic persistence of *P. aeruginosa*. Cumulatively, this live imaging analysis revealed that persistence likely occurs as bacterial clusters and within an intra-macrophage niche.

### High resolution imaging of the intracellular niche of persistent bacteria

As described above, live imaging revealed the presence of persistent *P. aeruginosa* bacteria, both inside and outside macrophages at 24 and 48 hpi (Figures 2A and 2G), but likely not within neutrophils (Figures 2F and 2G). Since *P. aeruginosa* can also enter and reside within non-phagocytic cells ^30-32^, we investigated their presence within non-phagocytic cells at the infection site. To this aim, we used Tg*(rcn3:Gal4/UAS:mCherry)* transgenic embryo harboring red fluorescent mesenchymal cells. Intracellular bacteria (individual or clusters) were clearly visualized at 24 and 48 hpi (Figure S6), demonstrating that *P. aeruginosa* also uses this niche *in vivo*, supporting pioneer work carried out with mice corneal infections ^9,32^.

We used electron microscopy (EM) to highlight the ultrastructural context of persistent bacteria in the tail fin of 24 or 48 hpi fixed infected Tg*(mfap4::mCherry-F)* embryos (Figure 3). Bacteria were largely seen inside cells, either isolated (Figure 3A) or in clusters (Figure 3B). A detailed analysis indicated that some bacteria were localized in vacuoles, as shown by the shape of clusters and/or the visualization of a vacuolar membrane, whereas other appeared in the cytoplasm (Figures 3A and 3B). Some cells with intracellular bacterial cluster appeared damaged (Figure 3C, left panel) and bacteria could be visualized in apparently cell remnants or in the extracellular space (Figure 3C, right panels). Since identifying the host-cell type only by ultrastructural criteria was challenging, we also performed correlative Light-Electron Microscopy (CLEM) (Figures 3D and 3E). A single large cluster of bacteria inside a macrophage was found by light microscopy (Figure 3D). The same embryo was processed for EM and serial lateral sections were prepared and imaged by array tomography (Figure 3E). The cluster showed a high number of bacteria inside a cell identified as a macrophage by the corresponding fluorescence image.

**Figure 3.**
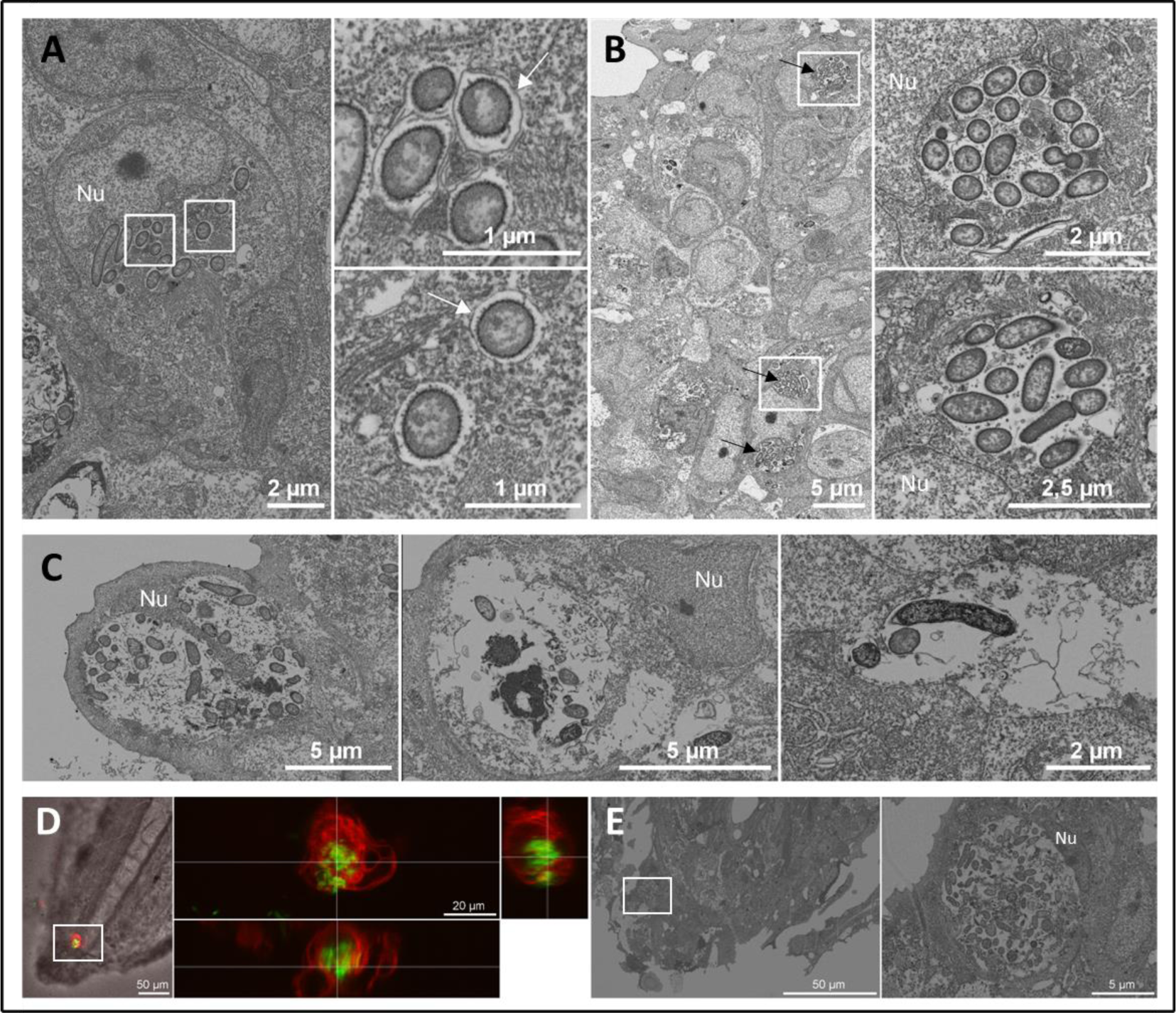
Electron micrographs of persistent *P. aeruginosa* B6513 in zebrafish. Representative images were acquired from a thin section of infected embryos at 24 hpi (D-E) or 48 hpi (A-C). (A) High magnification images of isolated intracellular bacteria near the tail fin edge (pixel size 5 nm). Right panels are zoom of the white squares. Nu: cell nucleus. A membrane is visualized around some bacteria (white arrow). (B) High magnification images of clustered intracellular bacteria with vacuolar shape (black arrows) at the tail fin. Right panels are zoom of the white squares. (C) High magnification images of intracellular bacteria in a damaged cell (left panel), or bacteria in apparently cell remnants (middle panel) or in the extracellular space (right panel). (D) and (E) Correlative Light-electron Microscopy (CLEM). (D) A z-stack of a live zebrafish embryo was acquired. The left panel shows a single fluorescent plane overlaid with the central DIC image of the fish tail (macrophages are seen in red and bacteria in green). A single large cluster containing dozens bacteria inside a macrophage is shown (white square). The right panel shows the orthogonal projection of the portion of the stack containing the cluster. (E) After processing the same fish for EM, lateral serial thin sections were prepared and imaged by EM. One section was acquired at a pixel size of 25 nm. Left panel shows an overview of the tail region. The large cluster observed in light microscopy is shown in its physiological context in the left panel (white square). The portion of the image containing the cluster is enlarged on the right panel, showing a high number of bacteria, filling almost all the cytoplasm of the cell and leaving the nucleus intact on one side.

Taken together, live and electron microscopy analyses indicated that *P. aeruginosa* bacteria persisting at the infection site were largely intracellular, in either macrophages or non-phagocytic cells. Moreover, CLEM imaging allowed visualization of a large cluster of *P. aeruginosa* inside a macrophage. These findings support the uniqueness of the zebrafish model in deciphering how *P. aeruginosa* interacts with host cells during infection.

### Strains with a persistent profile in zebrafish survived inside cultured macrophages and formed biofilms *in vitro*

In our *in vivo* model, persistent *P. aeruginosa* were visualized as intramacrophage bacterial clusters at the zebrafish wound site (Figures 2 and 3). We addressed the *in vitro* features of the CF isolates by investigating their phenotypes in a macrophage cell line and in a biofilm assay. We first monitored the intra-macrophagic survival of the different clinical strains by performing a gentamicin protection assay in murine J774 cells infected at a multiplicity of infection (MOI) of 10, as done previously with PAO1 ^33,34^. After 1.5 h of internalization, quantification of intracellular bacteria revealed that strain A6520 was largely eliminated, while the number of C6490 and RP73 cells were only modestly reduced (Figure 4A). On the other hand, B6513 was not eliminated and survived inside macrophages for at least 1.5 h after phagocytosis (Figure 4A). Phagocytic rates of the different strains were not significantly different (Figure 4B). Time-lapse microscopy performed during 3 h after phagocytosis correlated with CFU measurements, with a striking clearance of strain A6520, but no clearance of macrophage cells, while B6513 was still residing within macrophages (Figure 4.C). The visualization of intra-vacuolar clusters with B6513 showed that this bacterium can reside in closed compartments within phagocytic cells (Figure 4D). We next evaluated the ability of the diverse strains to form biofilms. While A6520, B6513 and RP73 produced a pellicle covering the broth medium, C6490 failed to form such a biofilm at the air-liquid interface (Figure 4E).

**Figure 4.**
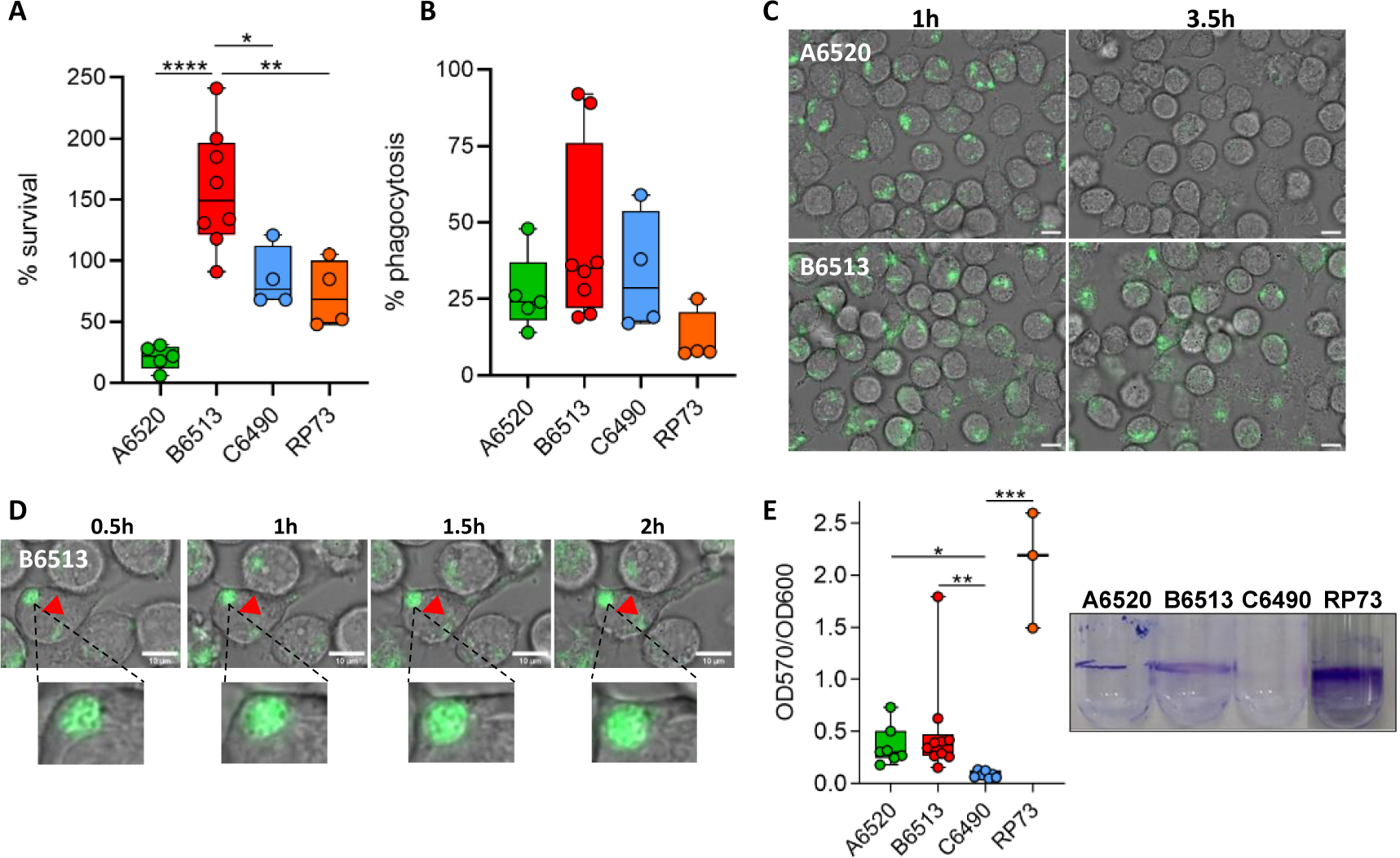
Persistent isolates are able to resist macrophages and form biofilm *in vitro*. (A) Bacterial survival within J774 macrophages (n=4 to 8). The number of intramacrophagic bacteria was determined 20 min (T0) or 2 h after gentamycin treatment (T1). Survival is expressed as the ratio between the numbers of bacteria recovered at T1 versus T0. One-way ANOVA: \*\*\*\**P*<0.0001; Tukey post-hoc test: \**P*<0.05, \*\**P*<0.01 and \*\*\*\**P*<0.0001. (B) Phagocytosis efficiency of each strain (n=4 to 8). The number of intracellular bacteria recovered at T0 was compared to the initial inoculum. (C) Representative initial and final images of time-lapse microscopy to follow the behavior of GFP^+^ intramacrophagic bacteria (isolates A6520 and B6513) for 2h30. Scale bar: 10 µm. (D) Time lapse imaging of an intra-vacuolar bacterial cluster (red arrowhead) visualized within a macrophage (a zoom is shown below). In (C) and (D), times after the start of gentamycin treatment are indicated. (E) Biofilm formation at 24h assessed by crystal violet (CV) assay on glass tubes (n=3 to 10). Quantification of the CV-labelled biofilm rings (a representative picture is shown on the right) is normalized with bacterial growth (OD_600nm_). Kruskal-Wallis test: \*\*\**P*<0.001; Dunn post-hoc test: \**P*<0.05, \*\**P*<0.01 and \*\*\**P*<0.001.

Overall, these data show that the two isolates recognized as persistent in the zebrafish embryo model combined the ability to resist killing by macrophages upon phagocytosis and the ability to form biofilms, whereas the two non-persistent isolates lacked one of these two traits.

### Persistent *P. aeruginosa* bacteria exhibit adaptive resistance to antibiotic treatment

Antibiotic adaptive resistance is a major issue in the treatment of chronic infections. We investigated antibiotic efficacy on embryos infected by the isolate B6513, by applying a 30 min treatment at 1.5, 24 and 48 hpi (Figure 5A). Three clinically-used antibiotics to fight *P. aeruginosa* infections were tested, namely tobramycin (an aminoglycoside), colistin (a polymyxin) and ciprofloxacin (a fluoroquinolone) ^35^. Antibiotics were used at 40 times the minimum inhibitory concentration (MIC; 40 µg/mL for tobramycin and 20 µg/mL for ciprofloxacin), except for colistin that was toxic for embryos and thus used at 2.5 x MIC (10 µg/mL). All drugs caused a sharp reduction in bacterial loads when applied soon after the initiation of infection (1.5 hpi), from approx. 20-fold (ciprofloxacin and tobramycin) to complete eradication (colistin) (Figure 5B). However, their efficacy was strongly reduced when used at 24 or 48 hpi, with colistin and ciprofloxacin decreasing CFU counts by less than 7-fold and tobramycin having no significant effect (Figure 5B). The ability of bacteria to withstand antibiotic challenges occurred at around the time of establishment of a persistent infection in the embryos, associated with intracellular and aggregated bacteria. This phenotype was not linked to the acquisition of mutations conferring resistance, since colonies isolated following ciprofloxacin challenge at 48 hpi remained fully sensitive to the antibiotic *in vitro*.

**Figure 5.**
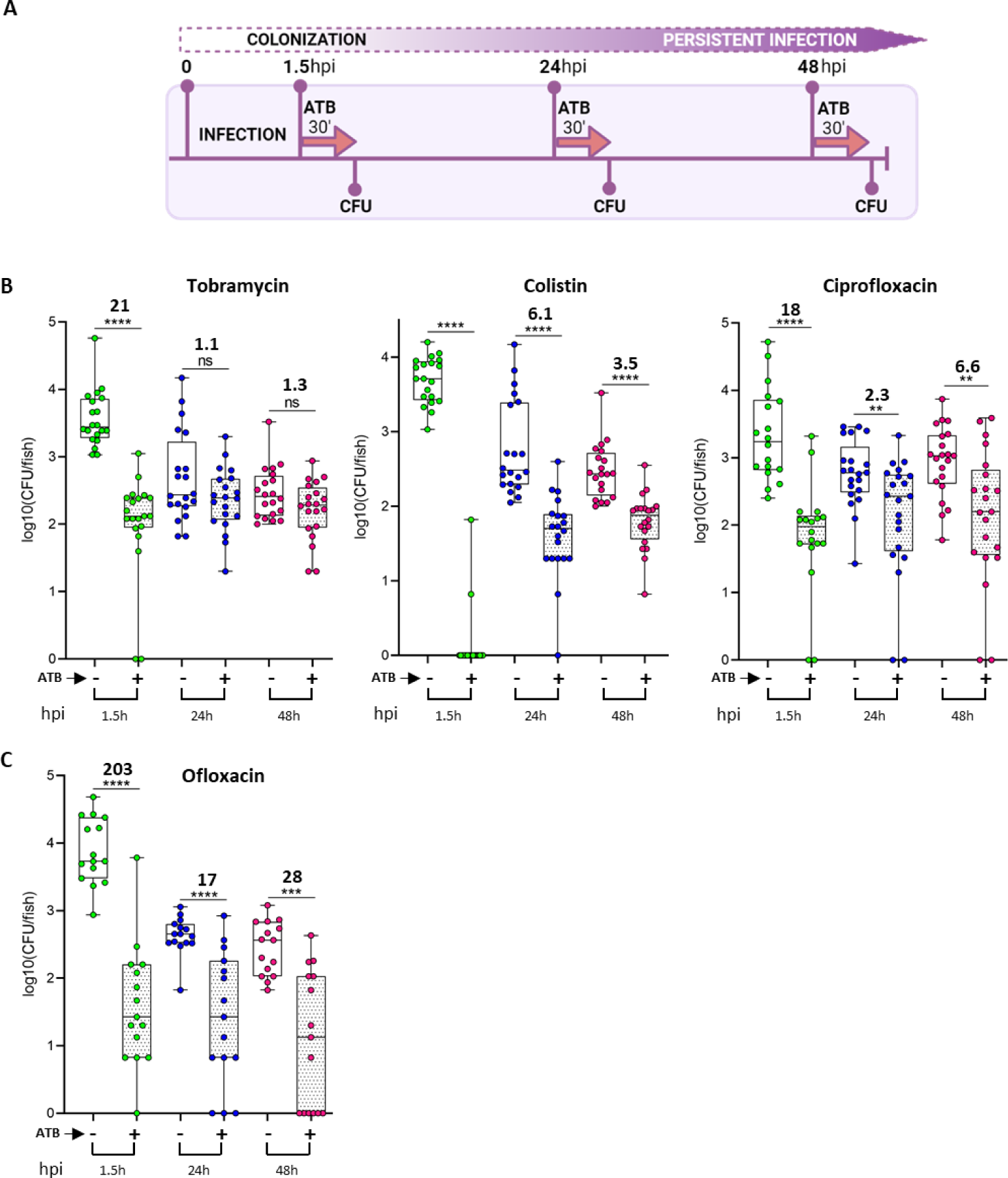
Antibiotics have a reduced efficacy on persistent *P. aeruginosa* in infected zebrafish. (A) Experimental procedure used to assess the efficacy of antibiotic treatments on infected embryos. ATB 30’: antibiotic treatment for 30 min. (B) Efficacy of various antibiotics on strain B6513 with respect to the time post-infection (n=3 to 4, 15 to 21 larvae). Embryos colonized for 1.5, 24 or 48 h were subjected to the indicated antibiotic challenge, or incubated in water for the control condition. Following this 30 min treatment, bacterial load per embryo was determined in both groups. Mann-Whitney test: \*\**P*<0.01, \*\*\**P*<0.001 and \*\*\*\**P*<0.0001. Ratios were calculated regarding the median of the data set; thus there is no values for colistin at t1.5 h as the median of the treated group is zero. (C) Efficacy of ofloxacin, an antibiotic known to enter eukaryotic cells with high efficiency, on strain B6513 (same analysis as in (B)).

During infection, *P. aeruginosa* may benefit from intracellular niches to escape antibiotics ^8,11^. Our microscopy analysis revealed the intracellular localization of persistent *P. aeruginosa*, which could *de facto* be protected from antibiotics (Figures 2A-C, Figure 3 and Figure S6). Antipseudomonal antibiotics differ in their ability to penetrate into eukaryotic cells, and their activity vary against intracellular *P. aeruginosa* in cultured phagocytes ^11^. To assess if the intracellular niche was responsible for the treatment failure in our model, we tested the *in vivo* activity of another fluoroquinolone, ofloxacin, that is known to be host cell permeable and to demonstrate better diffusion into human tissues than ciprofloxacin ^36^. Persistent bacteria at 24 and 48 hpi exhibited some tolerance towards ofloxacin (used at 40 x MIC, 20 µg/mL), when compared to bacteria treated very early after the start of infection. However, among the four antibiotics tested, ofloxacin was the most efficient molecule at the persistent stages (24 and 48 hpi), thus reinforcing the notion that eradication of intracellular persistent bacterial reservoirs requires administration of antibiotics able to reach these niches (Figure 5C). To complete this finding, we imaged Tg*(mfap4::mCherry-F)* embryos with persistent bacteria after treatment with ofloxacin and compared with images of non-treated embryos. The percentage of larvae with intra-macrophage bacteria dropped to 30% in ofloxacin-treated fishes, whereas it remained around 70% in non-treated controls.

Taken together, our results showed that the persistence of *P. aeruginosa* in zebrafish embryos is associated with an adaptive phenotype of resistance to antibiotics. Importantly, the observation that the most cell-permeant antibiotic was also the most effective one, suggests that reservoirs tolerant to other antibiotics are located within cells.

### Anti-biofilm compounds potentiate antibiotic efficacy on persistent bacteria

In our zebrafish model, persistent *P. aeruginosa* formed aggregates, which may display a higher tolerance to antibiotics than free-living bacteria ^7^. We hypothesized that anti-biofilm compounds would help to disassemble bacterial aggregates, thereby re-sensitizing persistent *P. aeruginosa* to antibiotics, potentially promoting their eradication with the assistance of the immune system. Among commercially available molecules with appealing activity against *P. aeruginosa* biofilm, we first used the human Atrial Natriuretic Peptide (hANP), an efficient biofilm-disperser *in vitro* that potentiates different antibiotics including tobramycin ^37^. Though devoid of antibacterial activity by itself, peptide hANP was able to reduce the formation of biofilms by strain B6513 *in vitro* (Figure S7). We then assessed its capacity to potentiate tobramycin and colistin treatments in infected embryos at 48 hpi, a time point at which bacteria were partly tolerant to antibiotics (Figure 6A). When applied alone, hANP modestly reduced bacterial loads by 2.5-fold, suggesting that it somehow assisted the innate immune system to eliminate bacteria. Strikingly, when used in combination with antibiotics, a more significant reduction of the pathogen burden was recorded i.e., 7-fold for tobramycin and 16-fold for colistin, when compared with the controls (Figure 6B). We performed similar experiments with a structurally different anti-biofilm compound, the fatty acid *cis-2-decenoic* acid (CDA). CDA, a molecule produced by *P. aeruginosa*, promotes biofilm dispersal and potentiates clearance of pre-established biofilms in combination with antibiotics ^38,39^. Moreover, CDA was proposed as a suitable molecule to target intra-macrophagic biofilms of pathogenic *Escherichia coli* ^40^. When used alone at 48 hpi, CDA had no detectable effects on persistent *P. aeruginosa* in zebrafish embryos, but when added in combination with colistin, bacterial loads per embryo decreased by 5-fold, thus increasing the antibiotic efficacy by >3-fold (Figure 6C).

**Figure 6.**
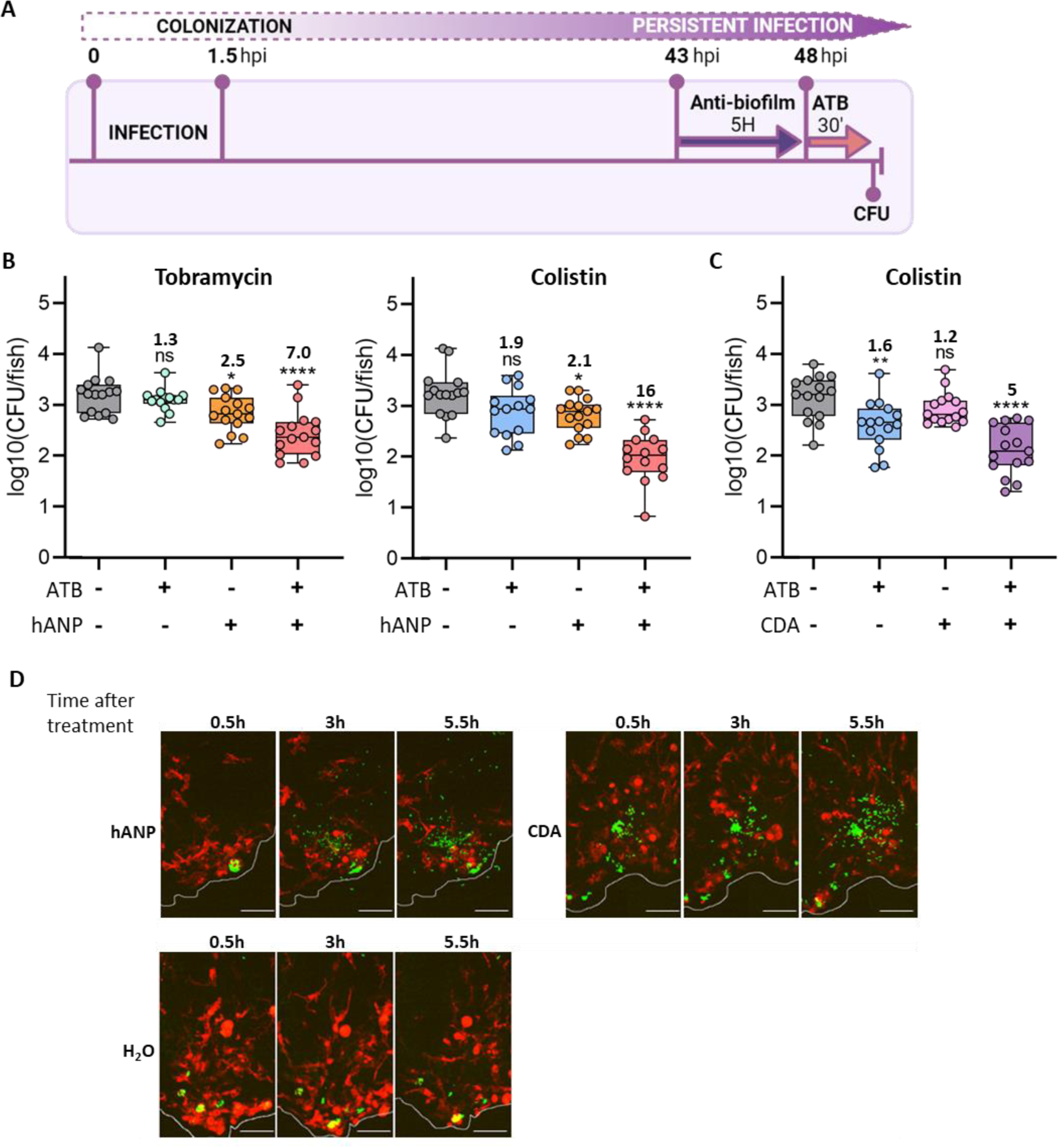
Treatment with anti-biofilm compounds can re-sensitize persistent *P. aeruginosa* bacteria to antibiotics in infected zebrafish. (A) Experimental procedure to assess the combined effect of anti-biofilm molecules and antibiotics. ATB 30’: antibiotic treatment for 30 min. (B) and (C) Antibiotic potentiation by 10 µM hANP (B) and 20 µM CDA (C) at 48 hpi (n=3, 14 to 15 larvae). Anti-biofilm compounds were added for 5 h to embryos infected by B6513-gfp strain, followed by the indicated antibiotic treatment, before CFU counting per embryo. One-way ANOVA: ****p<0.0001; Dunnett post-hoc test (comparison with the control condition): \**P*<0.05, \*\**P*<0.01 and \*\*\*\**P*<0.0001. Ratios were calculated regarding the median of the data set. (D) Maximal projections of confocal images, showing the effect of hANP or CDA following the treatment used in (B) and (C), without antibiotic treatment. Pictures of Tg(*mfap4:mCherry-F*) larvae infected by strain B6513-gfp were tan, from the beginning of the treatment, every 30 min for 5.5 h. A selection of images at 3 time points is shown. Scale bar: 40 µm.

We next took advantage of the optical transparency of embryos to visualize *in vivo* the effect of these anti-biofilm molecules on persistent bacteria. After a 5 h exposure to hANP or CDA, a reduction of the size of some bacterial clusters and concomitant increased number of isolated bacteria were observed, which was not the case in the H_2_O control conditions (Figures 6D and S8).

Cumulatively, these results are consistent with the notion that persistent *P. aeruginosa* bacteria present in aggregated structures, which are dispersible upon addition of anti-biofilm molecules, contribute to the adaptive antibiotic-resistant phenotype. Moreover, these findings validate the pertinence of our *in vivo* model to identify compounds that could potentiate the effects of conventional antibiotics.

## Discussion

Modeling bacterial chronic infection *in vivo* is essential to understand pathogenesis and evaluate treatment efficacy in a context that includes responsive host innate immunity and tissues. Here, we propose a novel *in vivo* model, complementary to mouse models, based on the infection of wounded zebrafish embryos with *P. aeruginosa* clinical strains. The zebrafish embryo model provides unique opportunities for intravital imaging and highlights the importance of an intracellular phase for establishing bacterial persistence in a vertebrate host. Moreover, this model recapitulates a major feature of chronicity, i.e. adaptive antibiotic resistance, and allows a simple screen for compounds that re-sensitize bacteria to antibiotics.

To establish this first model of persistent *P. aeruginosa* infection in zebrafish, we combined different parameters that have not been tested with clinical isolates to date: (i) parallel analysis of several *P. aeruginosa* isolates from CF patients, including a reference isolate known to establish a long-term infection in a murine model of chronic pneumonia ^24^, (ii) experiments over a period of up to 3 days after infection, and (iii) wound infection. The bacterial isolates could not be differentiated in this model based on the mortality rate of embryos since, in contrast to PAO1 strain, the mortality was very low or absent with the four isolates tested. However, the analysis over a 3 day period allowed us to clearly distinguish two classes of strains, those being persistent and those that were eliminated. Notably, the virulence and bacterial load profiles of clinical isolates in zebrafish perfectly matched the profiles observed in a mouse model of chronic wound infection, further validating the physiological relevance of our non-mammalian model.

The zebrafish embryo offers unique advantages to track bacterial infections in real time, and to study the interaction between bacteria and host cells. In keeping with previous studies using other infection routes, intravital microscopy revealed that infecting *P. aeruginosa* introduced at the tail injury were phagocytosed by both neutrophils and macrophages shortly after infection ^20,41^. While neutrophils were still present at the injury site at later time points, they rarely interacted with persistent *P. aeruginosa*, as opposed to macrophages. Unexpectedly, persistent bacteria were often visualized inside macrophages, which are long-living cells, indicating that *P. aeruginosa* displays strategies to resist macrophage killing and can use macrophages as a persistence niche. Such strategies are hallmarks of diverse intracellular pathogens, and were not expected for *P. aeruginosa*, which is considered an extracellular bacterium. This is however consistent with our findings highlighting an intramacrophage stage in cellular models and during *P. aeruginosa* acute infection ^20,33,34^. In addition, we recently reported that strain PAO1 can persist at low levels in microinjected larvae that overcome or do not develop an acute infection, and some persistent bacteria were also visualized inside macrophages ^42^. Here high-resolution imaging (EM) of infected zebrafish confirmed that persistent bacteria often reside inside host cells. Whereas the intracellular location of *P. aeruginosa* during infection was first reported three decades ago ^32^, its relevance *in vivo* has been a matter of debate and its role in persistence has not to date been addressed. Importantly, recent studies using intravital confocal imaging of infected mouse corneas and confocal imaging of human lung explants from CF patients support the intracellular location of *P. aeruginosa* during infection, as individual bacterial or small bacterial clusters ^10,43^. The nature of cells harboring *P. aeruginosa* was however not ascertained in these studies and, in this regard, the zebrafish model is a unique model, complementary to mice and human studies, that enables monitoring of the dynamics of intracellular *P. aeruginosa in vivo*. Intra-animal high resolution correlative microscopy (CLEM) in zebrafish allowed us to image for the first time an intramacrophage *P. aeruginosa* cluster. Moreover, real time imaging demonstrated that such clusters can last for more than 10 hours inside macrophages. In infected zebrafish, intracellular persistent bacteria were also visualized in non-phagocytic cells, supporting the importance of invasion of non-phagocytic cells during infection ^43,44^.

Our previous studies with PAO1 derived strains infecting cultured macrophages showed diverse outcomes, including intracellular bacteria that reside in vacuoles or in the cytoplasm ^33,42^. We observed here a vacuolar location for isolate B6513 upon infection of a macrophage cell line, and high-resolution EM imaging from infected zebrafish confirmed that individual or clustered persistent bacteria can reside inside host cell vacuoles *in vivo*. This is in line with the recent finding of a vacuolar persistent subpopulation of intracellular *P. aeruginosa* in cultured epithelial cells and the intravital imaging of colored puncta considered as vacuolar bacterial clusters during corneal infection ^43^. A vacuolar location should contribute to clustering of bacteria, which may retain a similar cluster structure in case of vacuolar rupture. Some clusters of persistent *P. aeruginosa* visualized in zebrafish had sizes above 10 µm, which is in line with aggregates found in CF lungs and chronic wounds ^45,46^ that are considered to represent non-attached biofilm-like structures ^47^. Of note, the bacterial aggregates observed in zebrafish could be dispersed, with a concomitant increase in isolated bacteria, upon addition of two different anti-biofilm molecules, hANP or CDA. Real time imaging of the dispersion of bacterial aggregates in a living animal demonstrates the unprecedented benefits of zebrafish embryo to follow the dynamic of bacterial clusters *in vivo*.

Persistent bacteria in the zebrafish model were refractory to antibiotic challenge, which mimics bacterial behavior during chronic infection in mammals and humans ^48^. Such adaptive antibiotic resistance of bacteria has been attributed to several features, including biofilm structures, which are well documented *in vitro* for *P. aeruginosa* ^7^. Here, the aggregated bacteria visualized *in vivo* appeared to contribute to adaptive resistance since anti-biofilm compounds could significantly re-sensitize persistent bacteria to antibiotic treatment. The bacterial intracellular stage is another feature allowing evasion of antibiotic challenge that recently attracted interest in the case of extracellular pathogens as *P. aeruginosa* ^8,49^. For example, the residence inside cultured lung epithelial cells reduced antibiotic efficiency ^12,50^ and intracellular *P. aeruginosa* in bladder epithelial cells promoted bacterial persistence and antibiotic tolerance in a mouse model of urinary tract infection ^13^. In our zebrafish model, the observation that the most efficient cell-permeant antibiotic, ofloxacin, demonstrates the highest efficacy supports the idea that the intracellular location provides a niche where persisting bacteria are less accessible to antibiotics. Ofloxacin, remains however only partially efficient against persistent bacteria, which may correlate with a poor efficacy against bacteria with a vacuolar location, which would further protect bacteria from antibiotics ^43^.

Finding new therapeutics to tackle chronic infections is critical and we provide a new platform to screen and evaluate *in vivo* the efficacy of treatments against persistent *P. aeruginosa*. Our model underscores the importance of strategies efficiently targeting the protective intracellular niche, which plays a crucial role in sustaining persistence within the host animal. This was not obvious for a pathogen such as *P. aeruginosa* which has been considered to be largely extracellular, but was consistent with the importance of an intramacrophage stage during infection for other major pathogens considered as extracellular pathogens, including *Acinetobacter baumannii* and *Streptococcus pneumonia* ^51,52^. Additionally our *in vivo* model reinforces the relevance of anti-biofilm strategies that disperse bacterial aggregates to re-sensitize *P. aeruginosa* to antibiotic treatments. This zebrafish model offers the possibility of insightful therapies, especially combination treatments, that could be transposed and refined in mammalian models, including man. Integrating studies in zebrafish and mouse models should greatly accelerate the identification of efficient treatments against *Pseudomonas*. It will also allow better understanding of the contribution of intracellular stages prior to the formation of biofilm-like structures in models of *P. aeruginosa* chronic infection.

## Materials and Methods

### Clinical strains used in the study and growth conditions

*P. aeruginosa* clinical strains were routinely grown at 37°C in lysogeny broth (LB) with shaking at 180 rpm. Clinical strains A6520, B6513 and C6490 were isolated at the teaching hospital of Besançon (France) from three individual cystic fibrosis patients aged 4, 1 and 17 years, respectively showing a transient (A6520, B6513) or chronic (C6490) lung colonization with *P. aeruginosa.* All these patients were treated with antibiotics to control the infection (clinical data not available). Genome sequencing was carried out to classify the strains. Sequence Types (STs) were determined according to the MLST scheme available at PubMLST (https://pubmlst.org). Strains A6520, B6513 and C6490 were found to belong to the same phylum as reference strain PAO1 ^53^, to harbor the exotoxin-encoding gene *exoS,* and to be genotypically distinct (ST274, ST27 and ST633, respectively). This Whole Genome Shotgun project has been deposited at DDBJ/ENA/GenBank under the accession JAWDIG000000000 (A6520), JAWDIH000000000 (B6513) and JAWDII000000000 (C6490). RP73 is a well described CF *P. aeruginosa* ^25^.

### Chemicals

All antibiotics were purchased from Sigma-Aldrich. Ciprofloxacin was dissolved at 25 mg/mL in 0.1 M hydrochloric acid. Ofloxacin was dissolved at 20 mg/mL in 1 M NaOH. Colistin sulfate salt and Tobramycin were dissolved at 8 mg/mL in ultrapure water. The human Atrial Natriuretic Peptide (hANP) was purchased from Tocris Bioscience (Bio-Techne) and was dissolved at 1 mg/mL in ultrapure water and stored at −20 °C. The cis-2-Decenoic acid (CDA) was purchased from Sigma-Aldrich and was dissolved at 5.8 mM in 100% DMSO and stored at -20°C.

### Minimum inhibitory concentration (MIC) assays

Susceptibility levels of strain B6513 to the fluoroquinolones ciprofloxacin (0.5 µg/mL) and ofloxacin (0.5 µg/mL), the polymyxin colistin (4 µg/mL), and the aminoglycoside tobramycin (1 µg/mL) were determined by the standard microdilution method in 96-well plates. Overnight cultures in LB broth were diluted in the same medium to an initial inoculum of OD_600nm_ = 0.1. MIC was defined as the lowest concentration that inhibited bacterial growth.

### Construction of stable fluorescent *P. aeruginosa* strains

Strains with chromosomally encoded GFP were obtained by triparental mating by using a recombinant integrative plasmid miniCTX carrying the PX2-GFP fusion ^54^ (from Ina Attree, Université Grenoble Alpes, France). Following overnight growth in LB broth with appropriate antibiotics, *E. coli* TOP10 carrying either helper plasmid pRK2013 ^34^ or miniCTX-PX2-GFP were mixed (20 µL of each culture) and deposited as drops onto the surface of LB plates for 2 h at 37°C. In parallel, the overnight cultures of *P. aeruginosa* strains were incubated at 42°C. 20 µL fractions of these stationary phase cultures were then individually added to the *E. coli* dry drops and left at 37°C for 5 h. The bacterial spots were then scrapped off and directly plated on LB plates containing Irgasan (25 µg/mL) and tetracycline (from 50 to 200 µg/mL) to select *P. aeruginosa* transconjugants. After 18 h at 37°C, several colonies were streaked out on LB medium to verify the stable GFP expression, and an individual GFP^+^ colony was cultured in liquid LB to obtain a glycerol stock.

### Zebrafish lines

Wild-type AB and Golden lines or AB/Golden mixed genetic background zebrafish were used for survival and CFU experiments. For live imaging, lines carrying fluorescent macrophages Tg(*mfap4:mCherry-F*)ump6TG ^55^, neutrophils Tg(*LysC:dsRed*)nz50 ^56^ or mesenchymal cells Tg(*rcn3:Gal4/UAS:mCherry*) ^57,58^ were used. Fish maintenance, staging and husbandry were performed as described ^59^. Eggs were obtained by natural spawning, collected in petri dishes and incubated at 28°C in fish water composed of 60 μg/mL sea salts (Instant Ocean) in distilled water supplemented with 0.4 mM NaOH and 0.1% methylene blue. Embryos and larvae were staged according to ^60^. For experiments, larvae were used at 2-day post-fertilization (dpf) until 5 dpf.

### Infection by immersion of injured zebrafish embryos

The protocol was essentially as described earlier ^23^, with few modifications. Overnight bacterial cultures were diluted at 1:20 in fresh LB broth and incubated until the OD_600nm_ reached approx. 0.8-1. Cultures were centrifuged at 4000 rpm for 10 min and re-suspended in fish water at approx. 10^7^ bacterial/mL. The bacterial load was determined by subsequent plating onto LB agar after dilution into phosphate-buffered saline (PBS). 2 dpf embryos, previously dechorionated, were anesthetized with tricaine (300 µg/mL, Sigma-Aldrich) and injured at the tail fin using 25-gauge needles under a stereomicroscope (Motic). Embryos were distributed into a 6-well plate containing 4 mL of bacterial suspension (or fish water as a control) immediately after the injury, and incubated at 28°C for 1.5 h. Two washes in fish water were subsequently performed (30 min in 10 mL and a few minutes in 4 mL, respectively) to eliminate bacteria in the bath. Finally, larvae were transferred individually into a 96-well plate (survival experiment) or a 24-well plate (CFU measurement and microscopy) containing fish water, and incubated using a light/dark cycle at 28°C. For survival curves, death was determined based on the absence of heart beat after visual inspection.

### Bacterial load measurement in infected embryos

Before CFU quantification, infected embryos were washed for a few minutes in 4 mL of fish water and subsequently crushed individually using a pestle in 100 µL of PBS. Then 100 µL of Triton X-100 (1% final concentration) were added for 10 min to liberate bacteria from residual cells/tissues. Following lysis, several dilutions in PBS were spotted on LB agar plates and incubated approx. 18 h at 37°C. Only GFP^+^ colonies were considered for counting. For CFU measurement following treatment, embryos were washed before the treatment and not prior the crushing (see below).

### Embryo imaging

After reaching the 50% epiboly stage, embryos were maintained in fish water supplemented with the melanization inhibitor Phenylthiourea (PTU) to prevent pigmentation. Before imaging, larvae were anesthetized with 300 µg/mL tricaine and placed in 35 mm glass-bottom dishes (FluoroDish, World Precision Instruments) and immobilized with 1% low-melting-point agarose (Sigma-Aldrich), covered with fish water after solidification. Zebrafish larvae were imaged at indicated hpi using the ANDOR CSU-W1 confocal spinning disk on an inverted NIKON microscope (Ti Eclipse) with ANDOR Zyla 4.2 sCMOS camera (40x water/NA 1.15 objective). Exposure time for all channels (green, red and DIC) was set at 50 ms. The 3D files generated by multi-scan acquisitions were processed using Image J and compressed into maximum intensity projections. Brightness and contrast were adjusted for better visualization. Deconvolution of images was done to improve resolution with Huygens (SVI).

### Quantification of macrophage recruitment and bacterial volumes

As a proxy to quantify macrophage recruitment at the injury site following spinning disk confocal imaging, the area formed by positive pixels (mCherry^+^) was measured at the median plan of each stack. To that aim, using Fiji, a threshold of positivity was applied using the ‘’IsoData’’ method. 3D reconstructions and volume quantification of GFP^+^ events were performed using the Imaris ‘’surface’’ tool.

### Treatment of infected embryos with antibiotics and anti-biofilm molecules

Prior any treatment, embryos were systematically washed for a few minutes in 4 ml of zebrafish water. Infected larvae were incubated with 40 x MIC of antibiotics, except for colistin (2.5 x MIC). Depending of the antibiotic, controls were done with fish water alone or supplemented with NaOH or HCl at the proper concentration. At least 5 embryos of the same condition were treated together for 30 min in 1 mL at room temperature. For CDA (20 µM) and hANP (10 µM), treatments were applied individually for 5 h at 43 hpi in 200 µL at 28°C. Control conditions were fish water alone (hANP) or supplemented with 0.34% DMSO (CDA). Independent experiments were systemically performed 3 times, with at least 5 larvae per condition (i.e minimum 15 embryos in total).

### Cutaneous infection in mice

Animal experiments were performed in accordance with the Canadian Council on Animal Care (CCAC) guidelines and were approved by the University of British Columbia Animal Care Committee protocol (A23-0030). Mice used in this study were outbred CD-1 mice (female). All animals were purchased from Charles River Laboratories, Inc. (Wilmington, MA, United States) and were 7-8 weeks of age at the time of experiments. Mice weighed 25 ± 2 g at the experimental start point and standard animal husbandry protocols were employed.

We tested the virulence of *P. aeruginosa* clinical isolates (A6520, B6513, and C6490) and the Liverpool Epidemic Strain LESB58 in a nuanced model of cutaneous high-density infection as previously described ^16^ with minor modifications. All strains were sub-cultured at 37°C with shaking (250 rpm) to an OD_600nm_ = 1.0 in LB. Cells were washed twice with sterile phosphate buffered saline (PBS) and resuspended to a final OD_600nm_ = 0.5 or 1.0 for clinical isolates or LESB58 strains, respectively. Strains were used to form high-density abscess infections to model invasive or chronic infections. Abscesses were formed by injection of 50 μL of bacteria on the left dorsum of mice for 3 or 7 days. Disease progression and overall welfare of animals was monitored daily up to day three, and weekly thereafter. At experimental endpoint, animals were euthanized using carbon dioxide followed by cervical dislocation, and abscess lesion size was measured using a caliper. Abscesses were harvested in PBS and homogenized using a Mini-Beadbeater (BioSpec Products, Bartlesville, OK, United States) for bacterial enumeration on LB. Two independent experiments containing 2-4 biological replicates each were performed.

### Electron microscopy

Samples were chemically fixed using 2.5% glutaraldehyde (Electron Microscopy Sciences # 16216) in fish water. After fixation, embryos were stored at 4°C in the fixation solution until subsequent processing. The procedure for embedding in resin was adapted from a previous method ^61^. All the procedure, except the overnight incubation in uranyl acetate, was performed using a Pelco Biowave® PRO+ Microwave processing systems (TED Pella) following the program indicated in Table S1. Samples were post-fixed with 2% osmium tetroxide (O_s_O_4_) in 0.1M cacodylate buffer pH 7.4 containing 5mM CaCl_2_, immediately followed by 1.5% K-ferrocyanide in the same buffer. After washing, samples were treated with 1% thiocarbohydrazide at 60°C, washed with distilled water before a second incubation in 2% OsO_4._ After washing, samples were then incubated overnight in 1% uranyl acetate at 4°C. Samples were next heated at 40°C and further processed in the microwave (Table S1), washed and incubated in lead aspartate pre-heated at 50°C. Dehydration was performed with growing concentrations of acetonitrile. Samples were then impregnated in Epon Hard+™ resin, and polymerized 48 h at 60°C. All chemicals were from EMS.

Thin serial sections were made using an UCT ultramicrotome (Leica) equipped with a Jumbo ultra 35° diamond knife (Diatome). Section ribbons were collected on silicon wafers (Ted Pella) with the help of an ASH2 manipulator (RMC Boeckler). Sections were imaged with a Zeiss Gemini 360 scanning electron microscope on the MRI EM4B platform under high vacuum at 1.5 kV. Final images were acquired using the Sense BSD detector (Zeiss) at a working distance between 3.5 and 4 mm. Mosaics were acquired with a pixel size of 5 nm and a dwell time of 3.2 µs.

### Macrophage infection and quantification of intracellular bacteria

J774 cells (murine macrophage cell line) were maintained at 37°C in 5% CO_2_ in Dulbecco’s modified Eagle medium (DMEM, Gibco) supplemented with 10% fetal bovine serum (FBS, Gibco). The infection of J774 macrophages by *P. aeruginosa* was carried out essentially as described previously ^34^. J774 macrophages (5x10^5^ cells/well) were infected by mid-log phase bacteria in PBS, at a MOI of approx. 10. Infection synchronization was done by a 5 min centrifugation at 1000 rpm of the 24-well plate, and bacterial phagocytosis was allowed to proceed for 25 min. Cells were then washed three times with sterile PBS and fresh DMEM medium supplemented with 400 µg/mL gentamicin was added, which was retained throughout the infection to kill non-phagocytosed bacteria. Macrophages were lysed after 20 min (T0) or 2 h (T1) of gentamicin treatment, by using 0.1% Triton X-100 and the number of viable bacteria was determined by subsequent plating onto LB agar plates. Survival rate at T1 was compared to the number of internalized bacteria at T0.

### Live microscopy on cultured macrophages

J774 macrophages were seeded in **glass bottom 8 wells μ-slide (Ibidi #80827)** in DMEM medium supplemented with 10 % FBS and infected with *P*. *aeruginosa* strains expressing GFP as described above. Imaging started after 30 min of phagocytosis, when the media was changed to DMEM supplemented with 400 μg/ml gentamicin until 3 hrs post phagocytosis. Cells were imaged using an inverted epifluorescence microscope (AxioObserver, Zeiss), equipped with an incubation chamber set-up at 37°C and 5% CO_2_ and a CoolSNAP HQ2 CCD camera (Photometrics). Time-lapse experiments were performed, by automatic acquisition of random fields using a 63X Apochromat objective (NA 1.4). The frequency of acquisition is indicated in figure legends. Image treatment and analysis were performed using Zen software (Zeiss).

### Biofilm formation

Biofilm formation was assessed in low magnesium medium in glass tubes 24 h at 30°C under static condition as described previously ^62^. After 24h, bacterial growth was measured by OD_600nm_ and tubes were carefully washed with water. The biofilm at the air-liquid interphase was stained using crystal violet (CV) 0.1% during 15 min at room temperature. After staining, tubes were washed with water and CV was extracted using acetic acid (30%) and quantified by measuring the OD_570nm_.

### *In vitro* efficiency of hANP

The flow cell system, which allows for continuous bacterial biofilm formation, is assembled, prepared and sterilized as described earlier ^63^. For studying the impact of hANP peptide on established strain B6513 biofilm, we used an established protocol ^37^. Briefly, bacterial cells from an over-night culture, were recovered by centrifugation (10 min, 7,500 rpm) and washed with sterile physiological water (0.9% NaCl). Each channel of the flow cell (1 mm x 4 mm x 40 mm, Bio centrum, DTU) was inoculated with 300 μL of bacterial suspension prepared at an optical density of OD_580nm_=0.1. Bacterial adhesion was allowed without any flow for 2 h at 37°C. After 2 h of adhesion, the LB medium was pumped with a flow rate of 2.5 mL/h at 37°C for 24 h. Next, the 24 h-old biofilm was exposed for 2 h to 300 µL of hANP (0.1 µM) or 300 µL of ultra-pure distilled water (control condition), added to each channel of the flow cell and without flow. Prior to image acquisition, biofilm cultures were then rinsed with LB medium using a 2.5 mL/h flow rate for 15 min. Finally, bacterial cells were marked with 5 µM of SYTO9 green-fluorescent dye (Invitrogen) and observed using confocal laser scanning microscopy (CLSM). CLSM observations of biofilms were performed using a Zeiss LSM710 microscope (Carl Zeiss Microscopy) using a x40 oil immersion objective. In order to capture the entire biofilm depth, images were taken every millimeter. For visualization and processing of three-dimensional (3D) images, the Zen 2.1 SP1 software (Carl Zeiss Microscopy) was used. Using the COMSTAT software (http://www.imageanalysis.dk/), quantitative analyses of image stacks were carried out ^64^.

### Statistical analysis

GraphPad Prism 8.3.0 was used to perform all statistical tests and create graphs. The indicated test used to analyze each dataset was chosen depending on the normality of the data. Multiple comparisons were done by one-way ANOVA or Kruskall-Wallis test, followed by Tukey’s or Dunn’s pairwise comparison, respectively. The Dunnett’s many-to-one comparison was used following a one-way ANOVA. Mann-Whitney U test was used to compare two groups.

### Ethics statement for zebrafish

All zebrafish experiments described in the present study were conducted at the University of Montpellier by following the 3rs -Replacement, Reduction and Refinement-principles according to the European Union guidelines for handling of laboratory animals (https://environment.ec.europa.eu/topics/chemicals/animals-science_en) and were approved by the “Direction Sanitaire et Vétérinaire de l’Hérault” and the “Comité d’Ethique pour l’utilisation d’animaux à des fins scientifiques” under reference CEEA-LR-B4-172-37. Breeding of adult fish adhered to the international guidelines specified by the EU Animal Protection Directive 2010/63/EU. All experiments were performed before the embryos free feeding stage (5 dpf) and did not fall under animal experimentation law according to the EU Animal Protection Directive 2010/63/EU. Embryos were euthanized using the anesthetic tricaine up to a lethal dose before bleach treatment. Embryo manipulation, handling, and euthanasia were performed by well-trained and authorized staff. Embryos were euthanized using an anesthetic overdose of buffered tricaine before bleach treatment.

## Acknowledgments

We thank M. Bour and K. Jeannot (French National Reference Centre for antimicrobial resistance, Besançon, France) for providing and sequencing the clinical strains A6520, B6513 and C6490, and A. Bragonzi (Milano, Italy) for providing the strain RP73. We thank the imaging facility MRI, member of the national infrastructure France-BioImaging infrastructure supported by the French National Research Agency (ANR-10-INBS-04, «Investments for the future») for photonic Microscopy at the MRI-DBS-UM platform and training by V. Diakou and E. Jublanc. We thank C. Gonzalez and V. Goulian for the Aquatic model facility ZEFIX from LPHI. We thank G. Lutfalla (LPHI, Montpellier, France) and P. Huber (Université Grenoble Alpes, CEA, France) for critical reading of the manuscript.

## Author contributions

Conceptualization: S.P., F.N. and A.B.B.P.

Methodology: S.P., F.N., R.E.W.H, P.P. and A.B.B.P.

Investigation: S.P., F.N., L.B., A.B., M.A.A., M.L., P.N.

Visualization: S.P., F.N., L.B., A.B., M.A.A., M.L.

Supervision: A.B.B.P.

Writing—original draft: S.P. and A.B.B.P.

Writing—review & editing: S.P., R.E.W.H, O.L., P.P. and A.B.B.P.

Funding acquisition: A.B.B.P.

## Data and materials availability

All data needed to evaluate the conclusions of this study are presented in the paper and/or the Supplementary Materials. Further information for resources and reagents should be directed to and will be fulfilled by A.B.B.P. at LPHI, CNRS, France. All unique/stable reagents generated in this study are available from A.B.B.P without restriction, except for *P. aeruginosa* clinical isolates and fluorescent derivatives that are available under materials transfer agreement (MTA).

## References

1. Qin, S.G., Xiao, W., Zhou, C.M., Pu, Q.Q., Deng, X., Lan, L.F., Liang, H.H., Song, X.R., and Wu, M. (2022). Pathogenesis, virulence factors, antibiotic resistance, interaction with host, technology advances and emerging therapeutics. Signal Transduct. Target. Ther. 7. 10.1038%2Fs41392-022-01056-1.

2. Garcia-Clemente, M., de la Rosa, D., Máiz, L., Girón, R., Blanco, M., Olveira, C., Canton, R., and Martinez-García, M.A. (2020). Impact of Pseudomonas aeruginosa infection on patients with chronic inflammatory airway diseases. J. Clin. Med. 9. 10.3390/jcm9123800.

3. Serra, R., Grande, R., Butrico, L., Rossi, A., Settimio, U.F., Caroleo, B., Amato, B., Gallelli, L., and de Franciscis, S. (2015). Chronic wound infections : the role of Pseudomonas aeruginosa and Staphylococcus aureus. Expert. Rev. Anti. Infect. Ther. 13, 605–613. 10.1586/14787210.2015.1023291.

4. Moradali, M.F., Ghods, S., and Rehm, B.H. A. (2017). Pseudomonas aeruginosa lifestyle: a paradigm for adaptation, survival, and persistence. Front. Cell. Infect. Microbiol. 7. 10.3389/fcimb.2017.00039.

5. Tacconelli, E., Carrara, E., Savoldi, A., Harbarth, S., Mendelson, M., Monnet, D.L., Pulcini, C., Kahlmeter, G., Kluytmans, J., Carmeli, Y., et al. (2018). Discovery, research, and development of new antibiotics: the WHO priority list of antibiotic-resistant bacteria and tuberculosis. Lancet Infect. Dis. 18, 318–327. 10.1016/S1473-3099(17)30753-3.

6. Brauner, A., Fridman, O., Gefen, O., and Balaban, N.Q. (2016). Distinguishing between resistance, tolerance and persistence to antibiotic treatment. Nat. Rev. Microbiol. 14, 320–330. 10.1038/nrmicro.2016.34.

7. La Rosa, R., Johansen, H.K., and Molin, S. (2022). Persistent bacterial infections, antibiotic treatment failure, and microbial adaptive evolution. Antibiotics-Basel 11. 10.3390/antibiotics11030419.

8. Kember, M., Grandy, S., Raudonis, R., and Cheng, Z.Y. (2022). Non-Canonical host intracellular niche links to new antimicrobial resistance mechanism. Pathogens 11. 10.3390/pathogens11020220.

9. Resko, Z.J., Suhi, R.F., Thota, A.V., and Kroken, A.R. (2024). Evidence for intracellular Pseudomonas aeruginosa. Journal of bacteriology. J. Bacteriol. 10.1128/jb.00109-24.

10. Malet, K., Faure, E., Adam, D., Donner, J., Liu, L., Pilon, S.J., Fraser, R., Jorth, P., Newman, D.K., Brochiero, E., et al. (2024). Intracellular Pseudomonas aeruginosa within the Airway Epithelium of Cystic Fibrosis Lung Tissues. Am. J. Respir. Crit. Care. Med. 10.1164/rccm.202308-1451oc.

11. Buyck, J.M., Tulkens, P.M., and Van Bambeke, F. (2013). Pharmacodynamic evaluation of the intracellular activity of antibiotics towards Pseudomonas aeruginosa PAO1 in a model of THP-1 human monocytes. Antimicrobial agents and chemotherapy 57, 2310–2318. 10.1128/aac.02609-12.

12. Garcia-Medina, R., Dunne, W.M., Singh, P.K., and Brody, S.L. (2005). Pseudomonas aeruginosa acquires biofilm-like properties within airway epithelial cells. Infect. Immun. 73, 8298–8305. 10.1128/iai.73.12.8298-8305.2005.

13. Penaranda, C., Chumbler, N.M., and Hung, D.T. (2021). Dual transcriptional analysis reveals adaptation of host and pathogen to intracellular survival of associated with urinary tract infection. PLoS pathog. 17. 10.1371/journal.ppat.1009534.

14. Lorenz, A., Pawar, V., Häussler, S., and Weiss, S. (2016). Insights into host-pathogen interactions from state-of-the-art animal models of respiratory infections. FEBS letters 590, 3941–3959. 10.1002/1873-3468.12454.

15. Reyne, N., McCarron, A., Cmielewski, P., Parsons, D., and Donnelley, M. (2023). To bead or not to bead: A review of lung infection models for cystic fibrosis. Front. Physiol. 14. 10.3389/fphys.2023.1104856.

16. Pletzer, D., Mansour, S.C., Wuerth, K., Rahanjam, N., and Hancock, R.E.W. (2017). New mouse model for chronic infections by gram-negative bacteria enabling the study of anti-infective efficacy and host-microbe interactions. mBio 8. 10.1128/mbio.00140-17.

17. Meeker, N.D., and Trede, N.S. (2008). Immunology and zebrafish: Spawning new models of human disease. Dev. Comp. immunol. 32, 745–757. 10.1016/j.dci.2007.11.011.

18. Pont, S., and Blanc-Potard, A.B. (2021). Zebrafish embryo infection model to investigate Pseudomonas aeruginosa interaction with innate immunity and validate new therapeutics. Front. Cell. Infect. Microbiol. 11, 745851. 10.3389/fcimb.2021.745851.

19. Torraca, V., and Mostowy, S. (2018). Zebrafish infection: from pathogenesis to cell biology. Trends. Cell. Biol. 28, 143–156. 10.1016/j.tcb.2017.10.002.

20. Moussouni, M., Berry, L., Sipka, T., Nguyen-Chi, M., and Blanc-Potard, A.B. (2021). Pseudomonas aeruginosa OprF plays a role in resistance to macrophage clearance during acute infection. Sci. Rep. 11, 359. 10.1038/s41598-020-79678-0.

21. Phennicie, R.T., Sullivan, M.J., Singer, J.T., Yoder, J.A., and Kim, C.H. (2010). Specific resistance to Pseudomonas aeruginosa infection in zebrafish is mediated by the cystic fibrosis transmembrane conductance regulator. Infect. Immun. 78, 4542–4550. 10.1128/iai.00302-10.

22. Kumar, S.S., Tandberg, J.I., Penesyan, A., Elbourne, L.D.H., Suarez-Bosche, N., Don, E., Skadberg, E., Fenaroli, F., Cole, N., Winther-Larsen, H. C et al. (2018). Dual transcriptomics of host-pathogen interaction of cystic fibrosis isolate Pseudomonas aeruginosa PASS1 with zebrafish. Front. Cell. Infect. Microbiol. 8, 406. 10.3389%2Ffcimb.2018.00406.

23. Nogaret, P., El Garah, F., and Blanc-Potard, A.B. (2021). A novel infection protocol in zebrafish embryo to assess Pseudomonas aeruginosa virulence and validate efficacy of a quorum sensing inhibitor in vivo. Pathogens 10. 10.3390/pathogens10040401.

24. Facchini, M., De Fino, I., Riva, C., and Bragonzi, A. (2014). Long term chronic Pseudomonas aeruginosa airway infection in Mice. Jove-J. Vis. Exp. 10.3791/51019.

25. Bianconi, I., Jeukens, J., Freschi, L., Alcalá-Franco, B., Facchini, M., Boyle, B., Molinaro, A., Kukavica-Ibrulj, I., Tümmler, B., Levesque, R. C., et al. (2015). Comparative genomics and biological characterization of sequential isolates from persistent airways infection. BMC genomics 16. 10.1186/s12864-015-2276-8.

26. Sousa, A.M., and Pereira, M.O. (2014). Diversification during infection development in cystic fibrosis lungs-A review. Pathogens 3, 680–703. 10.3390%2Fpathogens3030680.

27. Brannon, M.K., Davis, J.M., Mathias, J.R., Hall, C.J., Emerson, J.C., Crosier, P.S., Huttenlocher, A., Ramakrishnan, L., and Moskowitz, S.M. (2009). Pseudomonas aeruginosa Type III secretion system interacts with phagocytes to modulate systemic infection of zebrafish embryos. Cell. Microbiol. 11, 755–768. 10.1111/j.1462-5822.2009.01288.x.

28. Clatworthy, A.E., Lee, J.S., Leibman, M., Kostun, Z., Davidson, A.J., and Hung, D.T. (2009). Pseudomonas aeruginosa infection of zebrafish involves both host and pathogen determinants. Infect. Immun. 77, 1293–1303. 10.1128/iai.01181-08.

29. Cafora, M., Deflorian, G., Forti, F., Ferrari, L., Binelli, G., Briani, F., Ghisotti, D., and Pistocchi, A. (2019). Phage therapy against Pseudomonas aeruginosa infections in a cystic fibrosis zebrafish model. Sci. Rep. 9, 1527. 10.1038/s41598-018-37636-x.

30. Mittal, R., Grati, M., Gerring, R., Blackwelder, P., Yan, D., Li, J.D., and Liu, X.Z. (2014). In vitro interaction of Pseudomonas aeruginosa with human middle ear epithelial cells. PloS one 9, e91885. 10.1371/journal.pone.0091885.

31. Angus, A.A., Lee, A.A., Augustin, D.K., Lee, E.J., Evans, D.J., and Fleiszig, S.M. (2008). Pseudomonas aeruginosa induces membrane blebs in epithelial cells, which are utilized as a niche for intracellular replication and motility. Infect. Immun. 76, 1992–2001. 10.1128/iai.01221-07.

32. Fleiszig, S.M., Zaidi, T.S., Fletcher, E.L., Preston, M.J., and Pier, G.B. (1994). Pseudomonas aeruginosa invades corneal epithelial cells during experimental infection. Infect. Immun. 62, 3485–3493. 10.1128/iai.62.8.3485-3493.1994.

33. Garai, P., Berry, L., Moussouni, M., Bleves, S., and Blanc-Potard, A.B. (2019). Killing from the inside: Intracellular role of T3SS in the fate of Pseudomonas aeruginosa within macrophages revealed by mgtC and oprF mutants. PLoS Pathog. 15, e1007812. 10.1371/journal.ppat.1007812.

34. Belon, C., Soscia, C., Bernut, A., Laubier, A., Bleves, S., and Blanc-Potard, A.B. (2015). A macrophage subversion factor is shared by intracellular and extracellular pathogens. PLoS Pathog. 11, e1004969. 10.1371/journal.ppat.1004969.

35. Papadimitriou-Olivgeris, M., Jacot, D., and Guery, B. (2022). How to manage Pseudomonas aeruginosa infections. Adv. Exp. Med. Biol. 1386, 425–445. 10.1007/978-3-031-08491-1_16.

36. Volpe, D.A. (2004). Permeability classification of representative fluoroquinolones by a cell culture method. AAPS PharmSci 6. 10.1208%2Fps060213.

37. Louis, M., Clamens, T., Tahrioui, A., Desriac, F., Rodrigues, S., Rosay, T., Harmer, N., Diaz, S., Barreau, M., Racine, P.J., et al. (2022). Pseudomonas aeruginosa biofilm dispersion by the human atrial natriuretic peptide. Adv. Sci. 9, e2103262. 10.1002/advs.202103262.

38. Rahmani-Badi, A., Sepehr, S., Fallahi, H., and Heidari-Kesher, S. (2015). Dissection of the *cis*-2-decenoic acid signaling network in using microarray technique. Front. Microbiol. 6. 10.3389/fmicb.2015.00383.

39. Sepehr, S., Rahmani-Badi, A., and Babaie-Naiej, H. (2014). Unsaturated fatty acid, *cis*-2-Decenoic Acid, in combination with disinfectants or antibiotics removes pre-established biofilms formed by food-related bacteria. PloS one 9. 10.1371/journal.pone.0101677.

40. Prudent, V., Demarre, G., Vazeille, E., Wery, M., Quenech’Du, N., Ravet, A., Dauverd-Girault, J., van Dijk, E., Bringer, M.A., Descrimes, M., et al. (2021). The Crohn’s disease-related bacterial strain LF82 assembles biofilm-like communities to protect itself from phagolysosomal attack. Commun. Biol. 4. 10.1038/s42003-021-02161-7.

41. McCarthy, R.R., Mazon-Moya, M.J., Moscoso, J.A., Hao, Y., Lam, J.S., Bordi, C., Mostowy, S., and Filloux, A. (2017). Cyclic-di-GMP regulates lipopolysaccharide modification and contributes to Pseudomonas aeruginosa immune evasion. Nat. Microbiol. 2, 17027. 10.1038/nmicrobiol.2017.27.

42. Hajjar, H., Berry, L., Wu, Y.Z., Touqui, L., Vergunst, A.C., and Blanc-Potard, A.B. (2024). Contribution of intramacrophage stages to Pseudomonas aeruginosa infection outcome in zebrafish embryos: insights from mgtC and oprF mutants. Sci. Rep. 14. 10.1038/s41598-024-56725-8.

43. Kumar, N.G., Nieto, V., Kroken, A.R., Jedel, E., Grosser, M.R., Hallsten, M.E., Mettrucio, M.M.E., Yahr, T.L., Evans, D.J., and Fleiszig, S.M.J. (2022). Pseudomonas aeruginosa can diversify after host cell invasion to establish multiple intracellular niches. mBio. 13, e0274222. 10.1128/mbio.02742-22.

44. Newman, J.N., Floyd, R.V., and Fothergill, J.L. (2022). Invasion and diversity in Pseudomonas aeruginosa urinary tract infections. J. Med. Microbiol. 71. 10.1099/jmm.0.001458.

45. Bjarnsholt, T., Alhede, M., Alhede, M., Eickhardt-Sorensen, S.R., Moser, C., Kühl, M., Jensen, P.O., and Hoiby, N. (2013). The in vivo biofilm. Trends Microbiol. 21, 466–474. 10.1016/j.tim.2013.06.002.

46. Lichtenberg, M., Kirketerp-Moller, K., Kvich, L.A., Christensen, M.H., Fritz, B., Jakobsen, T.H., and Bjarnsholt, T. (2023). Single cells and bacterial biofilm populations in chronic wound infections. Apmis. 10.1111/apm.13344.

47. Kragh, K.N., Tolker-Nielsen, T., and Lichtenberg, M. (2023). The non-attached biofilm aggregate. Commun. Biol. 6. 10.1038/s42003-023-05281-4.

48. Pang, Z., Raudonis, R., Glick, B.R., Lin, T.J., and Cheng, Z.Y. (2019). Antibiotic resistance in Pseudomonas aeruginosa: mechanisms and alternative therapeutic strategies. Biotechnol. Adv. 37, 177–192. 10.1016/j.biotechadv.2018.11.013.

49. Lamberti, Y., and Surmann, K. (2021). The intracellular phase of extracellular respiratory tract bacterial pathogens and its role on pathogen-host interactions during infection. Curr. Opin. Infect. Dis. 34, 197–205. 10.1097/qco.0000000000000727.

50. Mirzaei, R., Mohammadzadeh, R., Sholeh, M., Karampoor, S., Abdi, M., Dogan, E., Moghadam, M.S., Kazemi, S., Jalalifar, S., Dalir, A., et al. (2020). The importance of intracellular bacterial biofilm in infectious diseases. Microb. Pathogenesis 147. 10.1016/j.micpath.2020.104393.

51. Sycz, G., Di Venanzio, G., Distel, J.S., Sartorio, M.G., Le, N.H., Scott, N.E., Beatty, W.L., and Feldman, M.F. (2021). Modern Acinetobacter baumannii clinical isolates replicate inside spacious vacuoles and egress from macrophages. PLoS Pathog. 17. 10.1371/journal.ppat.1009802.

52. Ercoli, G., Fernandes, V.E., Chung, W.Y., Wanford, J.J., Thomson, S., Bayliss, C.D., Straatman, K., Crocker, P.R., Dennison, A., Martinez-Pomares, et al. (2018). Intracellular replication of Streptococcus pneumoniae inside splenic macrophages serves as a reservoir for septicaemia. Nat. Microbiol. 3, 600–610. 10.1038/s41564-018-0147-1.

53. Holloway, B.W. (1955). Genetic recombination in Pseudomonas aeruginosa. J. Gen. Microbiol. 13, 572–581. 10.1099/00221287-13-3-572.

54. Pont, S., Fraikin, N., Caspar, Y., Van Melderen, L., Attrée, I., and Cretin, F. (2020). Bacterial behavior in human blood reveals complement evaders with some persister-like features. PLoS Pathog. 16, e1008893. 10.1371/journal.ppat.1008893.

55. Phan, Q.T., Sipka, T., Gonzalez, C., Levraud, J.P., Lutfalla, G., and Mai, N.C. (2018). Neutrophils use superoxide to control bacterial infection at a distance. PLoS Pathog. 14, e1007157. 10.1371/journal.ppat.1007157.

56. Hall, C., Flores, M.V., Storm, T., Crosier, K., and Crosier, P. (2007). The zebrafish lysozyme C promoter drives myeloid-specific expression in transgenic fish. Bmc. Dev. Biol. 7. 10.1186/1471-213x-7-42.

57. Ellis, K., Bagwell, J., and Bagnat, M. (2013). Notochord vacuoles are lysosome-related organelles that function in axis and spine morphogenesis. J. Cell. Biol. 200, 667–679. 10.1083/jcb.201212095.

58. Nguyen-Chi, M., Luz-Crawford, P., Balas, L., Sipka, T., Contreras-López, R., Barthelaix, A., Lutfalla, G., Durand, T., Jorgensen, C., and Djouad, F. (2020). Pro-resolving mediator protectin D1 promotes epimorphic regeneration by controlling immune cell function in vertebrates. Br. J. Pharmacol. 177, 4055–4073. 10.1111/bph.15156.

59. Begon-Pescia, C., Boireau, S., Boyer-Clavel, M., Lutfalla, G., and Nguyen-Chi, M. (2022). Preparing sequencing grade RNAs from a small number of FACS-sorted larvae macrophages isolated from enzyme free dissociated zebrafish larvae. MethodsX 9. 10.1016/j.mex.2022.101651.

60. Kimmel, C.B., Ballard, W.W., Kimmel, S.R., Ullmann, B., and Schilling, T.F. (1995). Stages of Embryonic development of the Zebrafish. Dev. Dyn. 203, 253–310. 10.1002/aja.1002030302.

61. Hua, Y.F., Laserstein, P., and Helmstaedter, M. (2015). Large-volume en-bloc staining for electron microscopy-based connectomics. Nat. Commun. 6. 10.1038/ncomms8923.

62. Moussouni, M., Nogaret, P., Garai, P., Ize, B., Vives, E., and Blanc-Potard, A.B. (2019). Activity of a synthetic peptide targeting MgtC on Pseudomonas aeruginosa intramacrophage survival and biofilm formation. Front. Cell. Infect. Microbiol. 9, 84. 10.3389%2Ffcimb.2019.00084.

63. Tolker-Nielsen, T., and Sternberg, C. (2011). Growing and analyzing biofilms in flow chambers. Curr. Protoc. Microbiol. Chapter 1, Unit 1B 2. 10.1002/9780471729259.mc01b02s21.

64. Heydorn, A., Nielsen, A.T., Hentzer, M., Sternberg, C., Givskov, M., Ersboll, B.K., and Molin, S. (2000). Quantification of biofilm structures by the novel computer program COMSTAT. Microbiol-Sgm 146, 2395–2407. 10.1099/00221287-146-10-2395.

