## Supplemental files for "Intracellular *Pseudomonas aeruginosa* persist and evade antibiotic treatment in a wound infection model"

Stéphane Pont *et al.*

**This PDF file includes:**

Figs. S1 to S8
Table S1

**Fig. S1.**

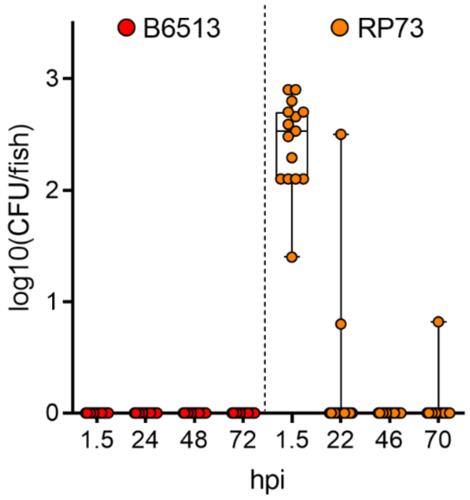


**Persistent isolates colonize the wound but not the skin of the embryos.** Uninjured larvae were immersed with GFP$^+$ persistent isolates B6513 and RP73, and were subsequently crushed and plated for CFU counting over approx. 72 hpi (n=3, 15 larvae).

**Fig. S2.**

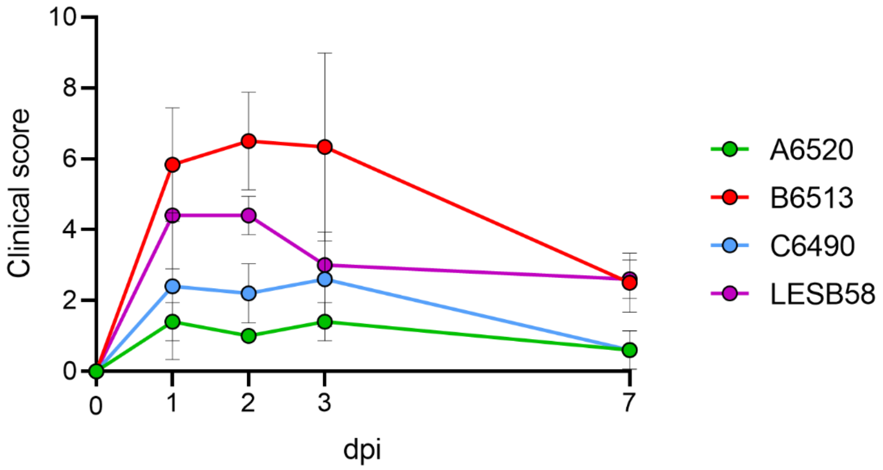


**Clinical scores of mice infected with the different *P. aeruginosa* clinical strains.** Clinical welfare of animals used in the murine model of high-density cutaneous infection was monitored using a standardized scoring system for assessing disease severity. Mice were monitored daily for the first three days post-infection (dpi), then weekly thereafter. Scores were assigned for a pre-determined battery of traits, including activity, hydration, pain, injection site, etc. and then summated for each animal. Strain B6513 did cause mortality of one animal at two dpi (out of 14 infected mice, data not shown), though the cause of death was not determined as total necropsy could not be performed prior to rigor mortis. Data are expressed as the mean clinical score for all animals in each treatment group +/- the standard error of the mean (SEM) ($n$ = 5-7).

**Fig. S3.**

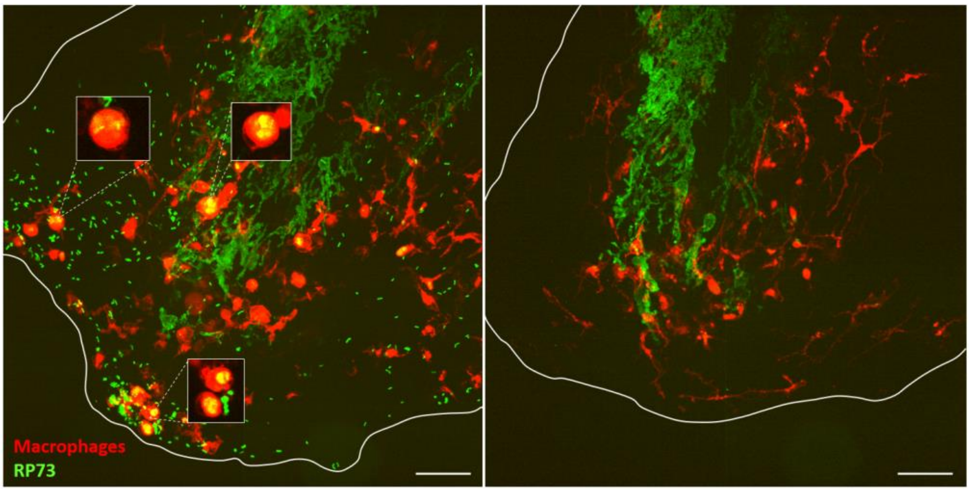


**Interaction of persistent RP73 strain with macrophages.** Left panel: representative maximal projection of confocal images, showing interactions between bacteria (green) and recruited macrophages (red) in Tg(*mfap4:mCherry-F*) larvae at 48 hpi. Boxed macrophages with intracellular *P. aeruginosa* were extracted from a single optical section. Right panel: non-infected embryo. Scale bar: 40 µm.

**Fig. S4.**

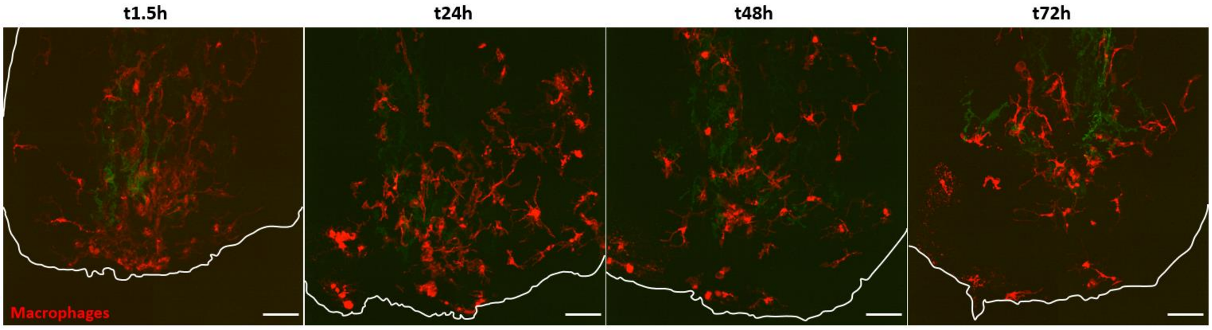


**Visualization of macrophages at the site of injury in absence of infection.** Maximal projections of confocal images, showing recruited macrophages (red) in Tg(*mfap4:mCherry-F*) larvae which were injured but not infected. Scale bar: 40 µm.

**Fig. S5.**

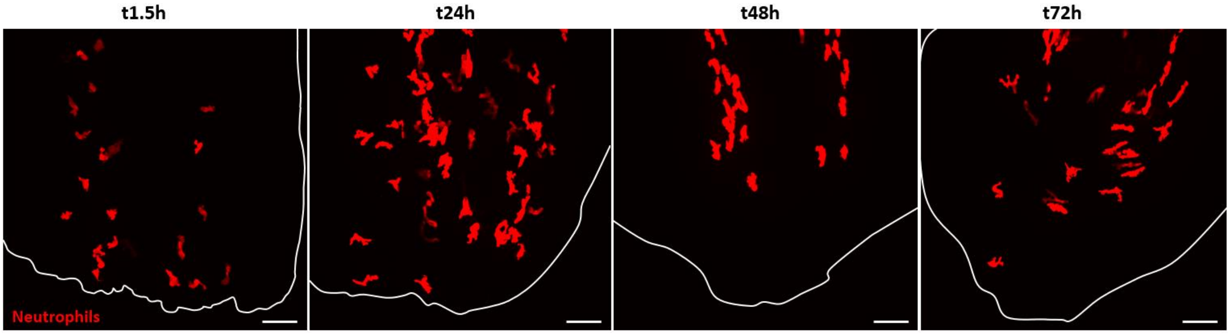


**Visualization of neutrophils at the site of injury in absence of infection.** Maximal projections of confocal images, showing recruited neutrophils (red) in Tg(*LysC*:*dsRed*) larvae which were injured but not infected. Scale bar: 40 µm.

**Fig. S6.**

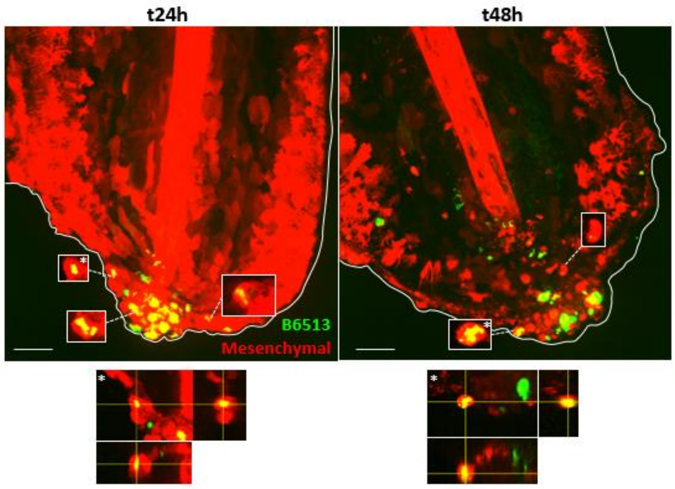


**Interaction of persistent *P. aeruginosa* B6513 with non-phagocytic cells**. Representative maximal projections of confocal images, showing interactions between bacteria (green) and mesenchymal cells (red) in Tg(*rcn3:Gal4/UAS:mCherry*) larvae at different time points. Boxed cells with intracellular *P. aeruginosa* were extracted from a single optical section. Below the images, orthogonal representations of the (*) boxed events, confirming that bacteria were intracellular. Scale bar: 40 µm. Note that pictures come from different embryos imaged for each indicated times.

**Fig. S7.**

A

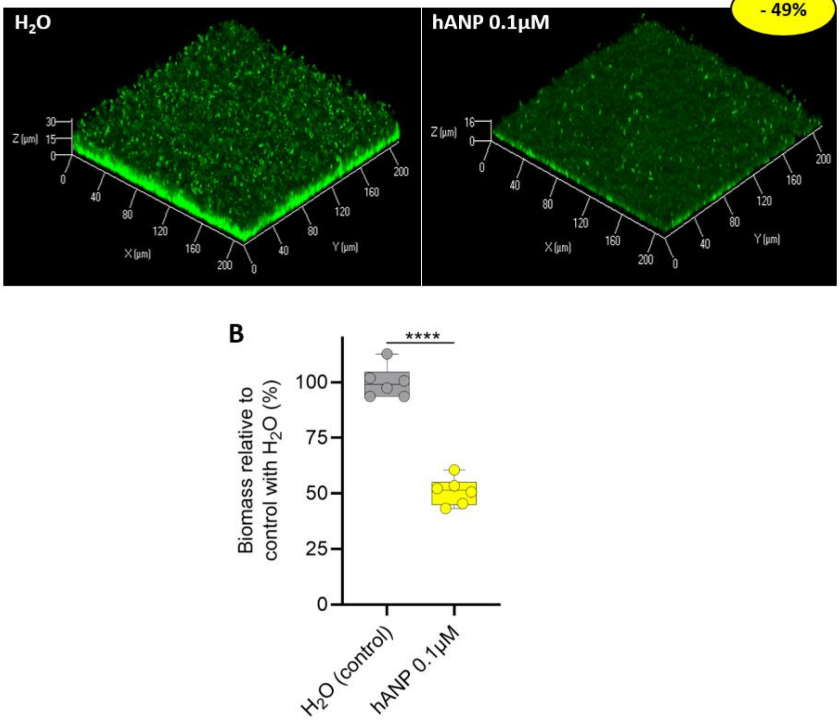


**Isolate B6513 responds to hANP *in vitro*. (A)** *In vitro* biofilms formed by the isolate B6513 for 24 h at 37°C in dynamic conditions were either untreated (left) or exposed to hANP at 0.1 µM (right) for 2 h, and were imaged by confocal microscopy following SYTO9 staining of bacterial cells. **(B)** COMSTAT analysis of biofilms imaged in (A), six views were extracted from two independent biological experiments (n=2). Student's t-test: ****p<0.0001.

**Fig. S8.**

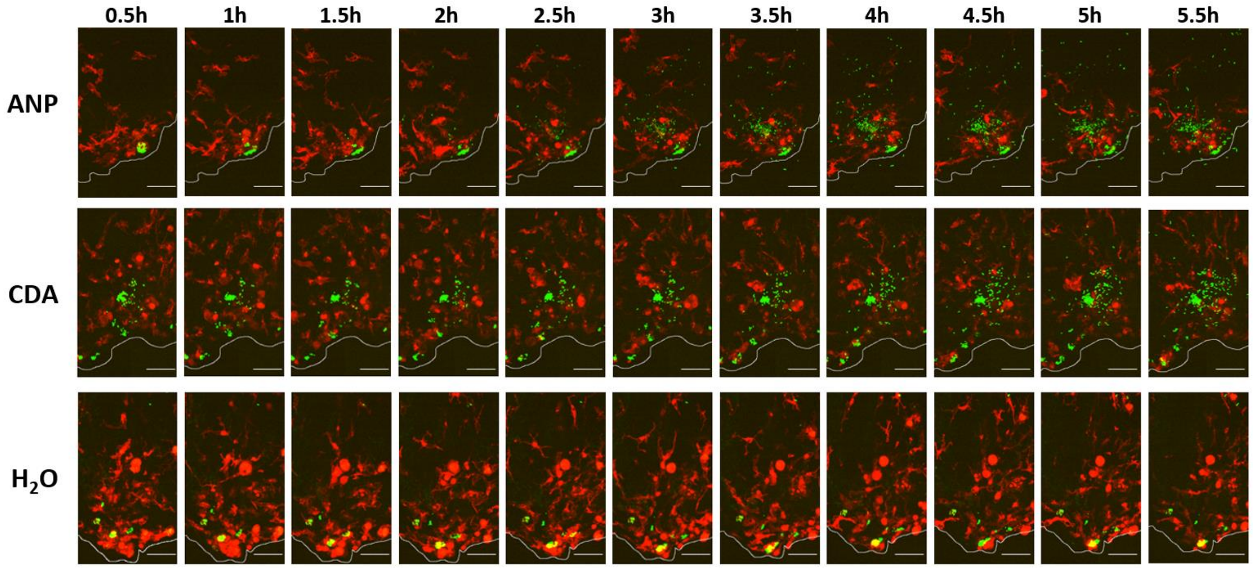


**Complete kinetic of the effect of anti-biofilm compounds.** Effect of hANP, CDA and $H_2O$ (control) shown in **Fig. 6**, with a shorter time frame between images (0.5 h compared to 2.5 h). Scale bar: 40 µm.

**Table S1.**

Glut = Glutaraldehyde; Ferro = K Ferrocyanide; UA = uranyl acetate

| Day 1 | step number | time h | time min | time sec | watts | vent time | vac time | set vac | load cooler T° | user prompt |
|---|---|---|---|---|---|---|---|---|---|---|
| Glut ON | 1 | 0 | 2 | 0 | 100 | 30 | 30 | 20 | 30 | 0 |
| Glut OFF | 2 | 0 | 2 | 0 | 0 | 30 | 30 | 20 | 30 | 0 |
| Glut ON | 3 | 0 | 2 | 0 | 100 | 30 | 30 | 20 | 30 | 0 |
| Glut OFF | 4 | 0 | 2 | 0 | 0 | 30 | 30 | 20 | 30 | 0 |
| Glut ON | 5 | 0 | 2 | 0 | 100 | 30 | 30 | 20 | 30 | 0 |
| Glut OFF | 6 | 0 | 2 | 0 | 0 | 30 | 30 | 20 | 30 | 0 |
| Glut ON | 7 | 0 | 2 | 0 | 100 | 30 | 30 | 20 | 30 | 0 |
| Buffer Rinse | 8 | 0 | 0 | 40 | 250 | 0 | 0 | 0 | 30 | 1 |
| Buffer Rinse | 9 | 0 | 0 | 40 | 250 | 0 | 0 | 0 | 30 | 1 |
| Buffer Rinse | 10 | 0 | 0 | 40 | 250 | 0 | 0 | 0 | 30 | 1 |
| OsO4 ON | 11 | 0 | 2 | 0 | 100 | 30 | 30 | 20 | 30 | 1 |
| OsO4 OFF | 12 | 0 | 2 | 0 | 0 | 30 | 30 | 20 | 30 | 0 |
| OsO4 ON | 13 | 0 | 2 | 0 | 100 | 30 | 30 | 20 | 30 | 0 |
| OsO4 OFF | 14 | 0 | 2 | 0 | 0 | 30 | 30 | 20 | 30 | 0 |
| OsO4 ON | 15 | 0 | 2 | 0 | 100 | 30 | 30 | 20 | 30 | 0 |
| OsO4 OFF | 16 | 0 | 2 | 0 | 0 | 30 | 30 | 20 | 30 | 0 |
| OsO4 ON | 17 | 0 | 2 | 0 | 100 | 30 | 30 | 20 | 30 | 0 |
| Ferro ON | 18 | 0 | 2 | 0 | 100 | 30 | 30 | 20 | 30 | 1 |
| Ferro OFF | 19 | 0 | 2 | 0 | 0 | 30 | 30 | 20 | 30 | 0 |
| Ferro ON | 20 | 0 | 2 | 0 | 100 | 30 | 30 | 20 | 30 | 0 |
| Ferro OFF | 21 | 0 | 2 | 0 | 0 | 30 | 30 | 20 | 30 | 0 |
| Ferro ON | 22 | 0 | 2 | 0 | 100 | 30 | 30 | 20 | 30 | 0 |
| Ferro OFF | 23 | 0 | 2 | 0 | 0 | 30 | 30 | 20 | 30 | 0 |
| Ferro ON | 24 | 0 | 2 | 0 | 100 | 30 | 30 | 20 | 30 | 0 |
| water Rinse | 25 | 0 | 0 | 40 | 250 | 0 | 0 | 0 | 30 | 1 |
| water Rinse | 26 | 0 | 0 | 40 | 250 | 0 | 0 | 0 | 30 | 1 |
| water Rinse | 27 | 0 | 0 | 40 | 250 | 0 | 0 | 0 | 30 | 1 |

| | | | | | | | | | | |
|---|---|---|---|---|---|---|---|---|---|---|
| TCH ON | 28 | 0 | 2 | 0 | 100 | 30 | 30 | 20 | 30 | 1 |
| TCH OFF | 29 | 0 | 2 | 0 | 0 | 30 | 30 | 20 | 30 | 0 |
| TCH ON | 30 | 0 | 2 | 0 | 100 | 30 | 30 | 20 | 30 | 0 |
| TCH OFF | 31 | 0 | 2 | 0 | 0 | 30 | 30 | 20 | 30 | 0 |
| TCH ON | 32 | 0 | 2 | 0 | 100 | 0 | 0 | 20 | 30 | 0 |
| TCH OFF | 33 | 0 | 2 | 0 | 0 | 30 | 30 | 20 | 30 | 0 |
| TCH ON | 34 | 0 | 2 | 0 | 100 | 30 | 30 | 20 | 30 | 0 |
| water Rinse | 35 | 0 | 0 | 40 | 250 | 0 | 0 | 0 | 30 | 1 |
| water Rinse | 36 | 0 | 0 | 40 | 250 | 0 | 0 | 0 | 30 | 1 |
| water Rinse | 36 | 0 | 0 | 40 | 250 | 0 | 0 | 0 | 30 | 1 |
| OsO4 2 ON | 37 | 0 | 2 | 0 | 100 | 30 | 30 | 20 | 30 | 1 |
| OsO4 2 OFF | 38 | 0 | 2 | 0 | 0 | 30 | 30 | 20 | 30 | 0 |
| OsO4 2 ON | 39 | 0 | 2 | 0 | 100 | 30 | 30 | 20 | 30 | 0 |
| OsO4 2 OFF | 40 | 0 | 2 | 0 | 0 | 30 | 30 | 20 | 30 | 0 |
| OsO4 2 ON | 41 | 0 | 2 | 0 | 100 | 30 | 30 | 20 | 30 | 0 |
| OsO4 2 OFF | 42 | 0 | 2 | 0 | 0 | 30 | 30 | 20 | 30 | 0 |
| OsO4 2 ON | 43 | 0 | 2 | 0 | 100 | 30 | 30 | 20 | 30 | 0 |
| water Rinse | 44 | 0 | 0 | 40 | 250 | 0 | 0 | 0 | 30 | 1 |
| water Rinse | 45 | 0 | 0 | 40 | 250 | 0 | 0 | 0 | 30 | 1 |

1

| | | | | | | | | | | |
|---|---|---|---|---|---|---|---|---|---|---|
| water Rinse | 46 | 0 | 0 | 40 | 250 | 0 | 0 | 0 | 30 | 1 |
| Overnight in 2% AcU 4°C then heat at 40°C (without washing) | | | | | | | | | | |
| **Day 2** | **step number** | **time h** | **time min** | **time sec** | **watts** | **vent time** | **vac time** | **set vac** | **load cooler T°** | **user prompt** |
| AcU 2 ON | 1 | 0 | 2 | 0 | 100 | 30 | 30 | 20 | 30 | 1 |
| AcU 2 OFF | 2 | 0 | 2 | 0 | 0 | 30 | 30 | 20 | 30 | 0 |
| AcU 2 ON | 3 | 0 | 2 | 0 | 100 | 30 | 30 | 20 | 30 | 0 |
| AcU 2 OFF | 4 | 0 | 2 | 0 | 0 | 30 | 30 | 20 | 30 | 0 |
| AcU 2 ON | 5 | 0 | 2 | 0 | 100 | 30 | 30 | 20 | 30 | 0 |
| AcU 2 OFF | 6 | 0 | 2 | 0 | 0 | 30 | 30 | 20 | 30 | 0 |
| AcU 2 ON | 7 | 0 | 2 | 0 | 100 | 30 | 30 | 20 | 30 | 0 |
| Rinse | 8 | 0 | 0 | 40 | 250 | 0 | 0 | 0 | 30 | 1 |
| Rinse | 8 | 0 | 0 | 40 | 250 | 0 | 0 | 0 | 30 | 1 |
| Rinse | 9 | 0 | 0 | 40 | 250 | 0 | 0 | 0 | 30 | 1 |
| Lead aspartate ON | 10 | 0 | 2 | 0 | 100 | 30 | 30 | 20 | 30 | 1 |
| Lead aspartate OF | 11 | 0 | 2 | 0 | 0 | 30 | 30 | 20 | 30 | 0 |
| Lead aspartate ON | 12 | 0 | 2 | 0 | 100 | 30 | 30 | 20 | 30 | 0 |
| Lead aspartate OF | 13 | 0 | 2 | 0 | 0 | 30 | 30 | 20 | 30 | 0 |
| Lead aspartate ON | 14 | 0 | 2 | 0 | 100 | 30 | 30 | 20 | 30 | 0 |
| Lead aspartate OF | 15 | 0 | 2 | 0 | 0 | 30 | 30 | 20 | 30 | 0 |
| Lead aspartate ON | 16 | 0 | 2 | 0 | 100 | 30 | 30 | 20 | 30 | 0 |
| Rinse | 17 | 0 | 0 | 40 | 250 | 0 | 0 | 0 | 30 | 1 |
| Rinse | 18 | 0 | 0 | 40 | 250 | 0 | 0 | 0 | 30 | 1 |
| Acetonitrile 50% | 19 | 0 | 0 | 40 | 250 | 0 | 0 | 0 | 30 | 1 |
| Acetonitrile 70% | 20 | 0 | 0 | 40 | 250 | 0 | 0 | 0 | 30 | 1 |
| Acetonitrile 80% | 21 | 0 | 0 | 40 | 250 | 0 | 0 | 0 | 30 | 1 |

| | | | | | | | | | | |
|---|---|---|---|---|---|---|---|---|---|---|
| Acetonitrile 90% | 22 | 0 | 0 | 40 | 250 | 0 | 0 | 0 | 30 | 1 |
| Acetonitrile 100% | 23 | 0 | 0 | 40 | 250 | 0 | 0 | 0 | 30 | 1 |
| Acetonitrile 100% | 24 | 0 | 0 | 40 | 250 | 0 | 0 | 0 | 30 | 1 |
| Acetonitrile 100% | 25 | 0 | 0 | 40 | 250 | 30 | 30 | 20 | 30 | 1 |
| Resin 50 | 28 | 0 | 3 | 0 | 150 | 30 | 30 | 20 | 30 | 1 |
| Resin 75 | 29 | 0 | 3 | 0 | 150 | 30 | 30 | 20 | 30 | 1 |
| Resin 100 | 30 | 0 | 3 | 0 | 150 | 30 | 30 | 20 | 30 | 1 |
| Resin 100 | 31 | 0 | 3 | 0 | 150 | 30 | 30 | 20 | 30 | 1 |
| Resin 100 | 32 | 0 | 3 | 0 | 150 | 30 | 30 | 20 | 30 | 1 |

**Detailed program of microwave processing used for resin embedding.**
